# KRAKEN: A provenance-tracked knowledge graph for multiomic and wellness research

**DOI:** 10.64898/2026.08.18.745544

**Authors:** Amy K. Glen, Drew Witherington, Trent Leslie, Andrew Baumgartner, Ashen Fernando, Bhargav Vemuri, Ornit Nahman, Gwênlyn Glusman, Leroy Hood, Lance Pflieger, Noa Rappaport

## Abstract

Existing general-purpose biomedical knowledge graphs tend to focus on disease mechanisms and drug repurposing, leaving multiomic and wellness-relevant content underrepresented. KRAKEN (Knowledge Research & Analysis Kit for Evidence Networks) addresses this gap by integrating existing graphs (including Translator KG Open, RTX-KG2, and ROBOKOP) with specialized sources such as RefMet, LIPID MAPS, NIH Common Data Elements, Polygenic Score Catalog, and derived wellness measures including biological age and biological BMI. The resulting graph spans ∼26M nodes and ∼121M edges across 62 entity types. KRAKEN adopts the Biolink Model as its semantic layer, ensuring compatibility with standardized resources emerging from the NIH NCATS Biomedical Data Translator program. A lightweight, modular build system rebuilds the full graph (including entity resolution), with peak memory consumption <48 GB, and supports flexible inclusion or exclusion of sources, allowing the user to scope the graph to a domain of interest. Built-in analytical tools include multi-hop reasoning, subgraph extraction, text, vector and hybrid entity search, and enrichment analyses, all accessible through an interactive web interface, a REST API, and a Model Context Protocol server, the last enabling direct consumption by agentic and LLM-based systems. KRAKEN is freely available at https://app.krakenkg.com.

**GRAPHICAL ABSTRACT:** 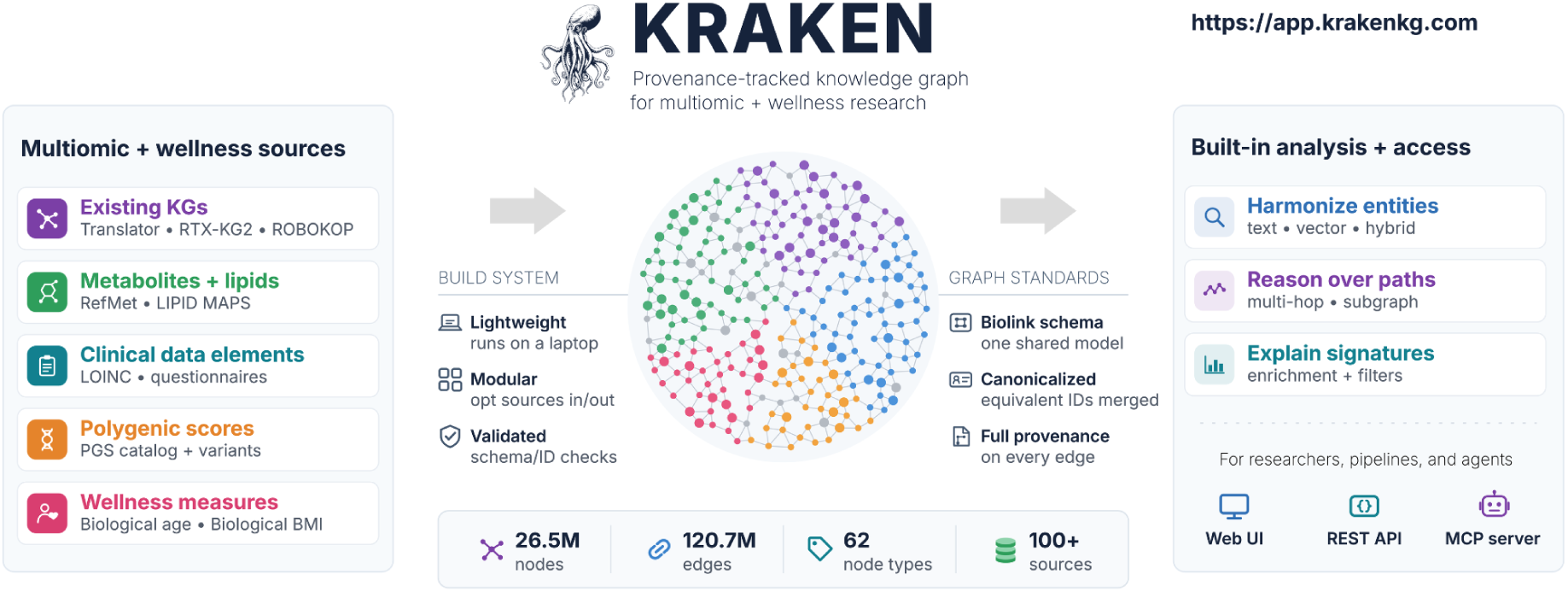

## INTRODUCTION

Efficient computational analysis and reasoning over biomedical knowledge requires that assertions scattered across curated databases, ontologies, and the published literature be brought into a single traversable structure. Existing knowledge graphs fall into two categories. The first harmonizes sources under a schema designed for the graph itself, as in Hetionet, which integrated 29 resources into a network whose path-based features prioritized drug repurposing candidates (1); PrimeKG, which organized 20 resources around disease nodes standardized to a single ontology (2); and SPOKE, which spans 27 million nodes and 53 million edges drawn from 41 databases (3). The second type fixes a shared semantic layer so that sources need not be reconciled twice: the NIH NCATS Biomedical Data Translator program (4) adopted the Biolink Model, which defines a hierarchy of entity categories and predicates together with a vocabulary for provenance and evidence (5), and the graphs built against it, among them RTX-KG2 (6) and ROBOKOP (7), are interoperable with each other and with the reasoning agents built over them. However, a shared semantic layer carries a cost. Mapping a source predicate onto the nearest category in a common vocabulary makes graphs interoperable while discarding distinctions the source drew, and reasoning over the harmonized result inherits those losses. The accessibility of these resources is often restricted by conflicting licenses among the sources they ingest. That constraint is a consequence of integrating broadly, and any graph assembled the same way inherits it. Across both categories, these knowledge graphs are designed to answer similar questions: which gene is implicated in a disease, which compound might be repurposed against it, and which mechanism connects the two.

Deep phenotyping cohorts pose a different level of complexity. Such studies profile multimodal datasets, often including untargeted metabolomics, plasma proteomics, gut microbiome composition, clinical chemistries, and polygenic risk on the same individuals, longitudinally and in people who are not sick (8); many also collect lipidomics and standardized questionnaire instruments (9). An analysis of such a cohort returns a mixed feature set, so its interpretation requires a graph in which most of those feature types are a resolvable entity. Metabolites and lipids appear largely through drug-target and disease-association edges rather than the functional and structural classifications that metabolomics platforms report, and functional interpretation of untargeted metabolomics remains limited by annotation coverage and by the fraction of measured features any pathway database represents (10, 11). Questionnaire instruments, standardized clinical data elements, polygenic scores, and derived multiomic measures such as biological age (BA) (12) and biological body mass index (bBMI) (13) are rarely represented as entities at all. The Clinical Knowledge Graph approached the problem from the assay side, embedding proteomics experiments in an open-source platform of close to 20 million nodes (14), which resolves the interpretive need for one assay type and leaves the cross-omic case open.

Frameworks exist for building custom graphs to fill such gaps: BioCypher (15) and KG-Hub (16) standardize the process of graph construction rather than any particular graph, as ORION does for ROBOKOP. Those frameworks address how a graph is assembled rather than what it contains, and they distribute graph artifacts, with or without a hosted query service. Retrieval at inference time reduces the rate at which language models produce unsupported statements (17, 18), and graph-derived context has been used to construct prompts for biomedical question answering (19). Yet the Cypher and TRAPI endpoints through which existing graphs are queried were specified before agentic consumption was a design target, and no standard defines what makes a graph ready for it. An additional class of tools uses language models to assemble graphs directly from text without mapping concepts to standard identifiers (20, 21). Those graphs suit exploration and hypothesis generation, and the absence of resolvable identifiers makes them unsuited to the harmonization and analysis tasks considered here. We therefore built KRAKEN to address five properties: 1) a semantic contract, so that a category or predicate name carries one meaning across every contributing source; 2) resolvable identity, so that an entity referred to by any source vocabulary returns the same canonical node; 3) evidence that travels with the assertion, so that a generated statement can be traced to a named source and an epistemic class rather than accepted on the model’s authority; 4) retrieval by more than one route, since identifier lookup, lexical search, semantic similarity, and structural traversal fail in different ways, together with the run-time discovery and bounded responses an agent needs to size a query before committing to it; and 5) reproducible construction, so that the graph can be regenerated as its sources change and a result attributed to a numbered release.

Herein, we describe KRAKEN, a hosted, Biolink-compliant knowledge graph of ∼26 million nodes and ∼121 million edges, developed within the ARPA-H PATH program and used both for harmonization, annotation, and analysis and as a substrate for agentic workflows. It integrates Translator ecosystem content with multiomic and wellness sources underrepresented in current community releases, among them RefMet, the LIPID MAPS Structure Database, the Polygenic Score Catalog, and NIH Common Data Elements. Polygenic scores, standardized clinical data elements, laboratory observation codes, functional metabolite classifications, and derived multiomic measures are not resolvable entities in Hetionet, PrimeKG, SPOKE, RTX-KG2, or ROBOKOP, while the analyses that generate them are the ones deep phenotyping cohorts produce. Representing them as entities under one semantic layer is what allows a mixed feature list to become a query. Like SPOKE, KRAKEN is served rather than distributed, through a web interface, a REST API, and a Model Context Protocol (MCP) server, with built-in tools for node discovery, multi-hop reasoning, subgraph extraction, and enrichment. In this paper, we present its sources and semantic layer, its build architecture and access infrastructure, the content and scale of the current release, and two use cases: mechanistic interpretation of a multiomic signature, and traversal from polygenic risk through correlated molecular mediators to candidate compounds. KRAKEN is freely available at https://app.krakenkg.com.

## MATERIALS AND METHODS

### Data Sources

KRAKEN integrates content from biomedical knowledge sources selected to support multiomic and wellness research. Directly ingested sources (**Table 1**) include distinctive resources with limited representation in general-purpose biomedical KGs (e.g., RefMet, LIPID MAPS, PGS Catalog) as well as established Translator ecosystem aggregator KGs (RTX-KG2, ROBOKOP, and Translator KG Open). In total, KRAKEN includes over 100 primary knowledge sources (including those ingested transitively through aggregator KGs), listed in Supplementary **Table S1**.

**Table 1.** Knowledge sources directly ingested into KRAKEN. Underlying knowledge sources ingested transitively through aggregator KGs are not listed here, but are included in Supplementary **Tables S1** and **S2**.

| Source | Version | Content type | License | Reference |
| --- | --- | --- | --- | --- |
| RefMet | Accessed 2026-08-07 | Metabolite nomenclature, cross-references, and classifications | CC BY-NC-ND | (22) |
| LIPID MAPS Structure Database | Accessed 2026-08-07 | Lipid nomenclature, cross-references, and classifications | CC BY 4.0 | (23) |
| Polygenic Score Catalog | Release 2026-07-29 | Polygenic score metadata and variant associations | EMBL-EBI Terms of Use* | (24) |
| NIH Common Data Elements | Accessed 2026-07-29 | Standardized questionnaire data element definitions for clinical research | ODC-ODbL | cde.nlm.nih.gov |
| LOINC | 2.82 | Laboratory test and clinical observation terminology | LOINC License† | (25) |
| UMLS | 2025AA | Cross-references between clinical terminologies (LOINC↔UMLS mappings) | UMLS Metathesaurus License‡ | (26) |
| NCBI Gene | Accessed 2026-08-31 | Gene nomenclature, cross-references, and summaries (selected taxa) | Unrestricted (NCBI data usage policy) | (27) |
| BA | – | Multi-omic biological age measure | – | (12) |
| bBMI | - | Multi-omic metabolic health measure | - | (13) |
| ISB Long COVID | Received 2026-09-17 | Long COVID associations with symptoms, proteins, and compounds | - | (28-31) |
| Microbiome KG | 2.1.0 | KG of relationships derived from supplemental tables in published microbiome papers | MIT License | (32) |
| Multiomics KG | 2.1.0 | KG of relationships derived from supplemental tables in published multiomics papers | MIT License | Goetz et al., in preparation |
| Clinical Trials KG | 4.1.7 | KG of intervention-condition relationships derived from clinical trial records | MIT License | (33) |
| Drug Approvals KG | 0.5.7 | KG of drug-condition approval relationships | MIT License | (33) |
| Translator KG Open | 2026_07_05 | Aggregator biomedical KG | MIT License | (4) § |
| RTX-KG2 | 2.10.2 | Aggregator biomedical KG | CC BY 4.0 | (6) |
| ROBOKOP | june2025 | Aggregator biomedical KG | MIT License | (7) |
| Babel | 2026jul22 | Identifier equivalence sets underlying Translator's SRI Node Normalizer | MIT License | (4) |

\* https://www.ebi.ac.uk/about/terms-of-use/
† https://loinc.org/kb/license
‡
https://www.nlm.nih.gov/research/umls/knowledge_sources/metathesaurus/release/license_agreement.html
§ Translator KG Open artifact downloaded from https://kgx-storage.ci.transltr.io/releases/translator_kg_open/

### Schema and Semantic Layer

KRAKEN uses the Biolink Model (5) for its schema and semantic layer. During queries of the hosted graph, node and edge types (e.g., SmallMolecule, interacts_with) are automatically traversed according to the Biolink category/predicate hierarchies, allowing queries of varying levels of abstraction to produce relevant answers. **Figure 1** shows an example node and edge from KRAKEN, demonstrating its schema. Nodes and edges carry standard Biolink-defined properties (described in more detail below) designed to support explainability. Source-specific data that does not necessarily fit into standard Biolink-defined properties but that may provide useful context for AI agents or humans is preserved in an ‘attributes’ property on nodes and edges, organized by contributing knowledge sources.

**Figure 1:**
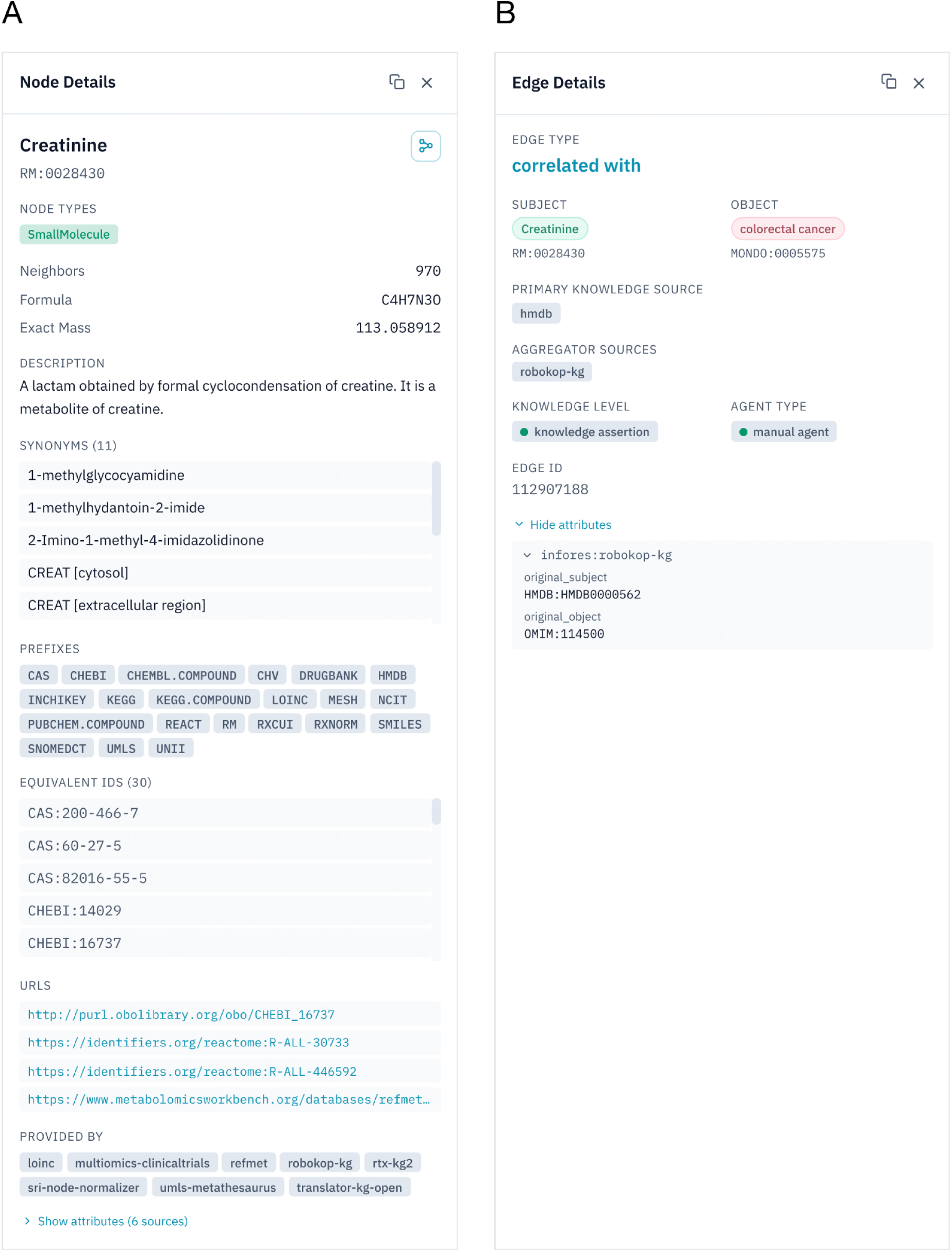
Example node and edge as they appear in the KRAKEN web user interface, demonstrating KRAKEN’s schema.

KRAKEN is represented in canonicalized form, in which equivalent identifiers from different ontologies/taxonomies (such as DOID:14330 and MONDO:0005180) are merged into one canonical node. Merged nodes retain identifiers from all contributing sources in an equivalent_ids property (**Figure 1A**), allowing queries against any source’s identifier to return the canonical entity. Edges with the same subject, predicate, object, primary knowledge source, and qualifiers are merged across aggregator sources to help avoid duplication of assertions from any shared underlying primary sources.

In full, every node has an identifier (id, a compact uniform resource identifier, or CURIE), one or more Biolink categories, the list of sources that provided it (provided_by), and the identifiers merged into it (equivalent_ids); optional node properties are name, description, synonyms, urls, taxon, publications, and, for chemicals, chemical_formula and exact_mass. Every edge has a subject and object (node identifiers), a Biolink predicate, a primary_knowledge_source, and a Biolink knowledge_level and agent_type; optional edge properties are qualifiers, aggregator_knowledge_source, supporting_data_sources, publications, and publications_info (excerpts from supporting publications, provided for text-mined edges). In addition, nodes and edges may both carry the ‘attributes’ property described above. Example node and edge records are provided in Supplementary Data S5.

### Build Architecture

KRAKEN is built by a lightweight, modularized Python-based pipeline that proceeds in two stages: per-source harmonization followed by cross-source integration. During harmonization, each source is mapped to KRAKEN’s Biolink-compliant schema to produce a subgraph in newline-delimited JSON (NDJSON) format. Controlled identifiers are normalized to standard CURIEs that follow Biolink conventions using the Normalizer module of the BioMapper Python package (https://github.com/Phenome-Health/biomapper2). During cross-source integration, entities are resolved using the equivalence information provided by the sources themselves: for Translator-aligned resources these identifiers derive from Translator normalization services (4), while for other resources (e.g., LIPID MAPS), equivalence relies on source-provided cross-references that include mappings not currently exposed by general-purpose normalization services. Equivalent nodes are merged into canonical entities and their corresponding edges are remapped accordingly, with provenance, publications, and attributes unioned across contributing sources. The build system uses a file streaming approach to keep peak memory consumption under 48 GB, with full builds taking less than 4 hours on modest hardware (MacBook Pro with Apple M4 chip), excluding source download time. Each harmonized source artifact and the final integrated graph are validated, with checks for orphan edges, Biolink compliance, and other integrity issues. The final graph is output as two NDJSON files (one for nodes, one for edges) and a summary metagraph in JSON format.

### Access Infrastructure

KRAKEN is accessible via three complementary interfaces, all of which expose the same underlying query capabilities and analytical tools, for different use cases:

1. An **interactive web user interface** for visual exploration, accessible at https://app.krakenkg.com

2. A **REST API** for programmatic access and incorporation into research pipelines, queryable at https://kestrel.krakenkg.com/api

3. An **MCP server** for direct consumption by LLMs and agentic systems, at https://kestrel.krakenkg.com/mcp

The MCP server uses Streamable HTTP transport and enables direct consumption by any MCP-compatible client (e.g., Claude Desktop, Cursor). KRAKEN’s MCP tools support multiple response modes: ‘slim’ for compact results, ‘full’ for complete detail, and ‘preview’ for result counts, enabling agents to iteratively refine queries before committing to full retrieval. Errors are returned as structured responses with status codes and messages, enabling agents to handle failures gracefully. Discovery tools (e.g., get_traversal_options) allow agents to enumerate available ranking presets, constraint fields, and operators at query time.

All three of KRAKEN’s interfaces are backed by KESTREL (Knowledge Engine for Structural, Text, and Representational Embedded Lookup), a Python-based query engine developed alongside KRAKEN. KESTREL loads the KRAKEN NDJSON files into read-only, memory-mapped structures: a key-value store (LMDB) holding node and edge records and an identifier-to-canonical-node lookup; compact numeric arrays encoding graph adjacency and the node and edge properties used for filtering; a full-text index (Tantivy) over node names, synonyms and descriptions; and a vector index (FAISS) over sentence embeddings of node names and synonyms. Text, vector and hybrid search identify matching nodes through the full-text and vector indexes, and multi-hop, query graph, subgraph, and enrichment tools identify nodes and edges through the adjacency and property arrays (both sets of tools are described under Analytical Tools); in every case the corresponding records are then retrieved from the key-value store. Biolink category and predicate hierarchies are resolved using the Biolink Model Toolkit (https://github.com/biolink/biolink-model-toolkit). The engine is exposed through a single REST API (FastAPI). The web interface is a browser application that calls this API, and the MCP server is a thin layer whose tools call the same endpoints, so all three interfaces return identical results from one implementation. Metadata about KRAKEN is accessible from all three interfaces, including its metagraph (https://kestrel.krakenkg.com/api/metagraph), source lists, and qualifier metadata. All KRAKEN API endpoints are documented at https://kestrel.krakenkg.com/api/docs.

### User interface

KRAKEN’s web interface provides interactive access to the analytical tools described in the following section. Here we show examples of two of those tools. **Figure 2** shows results of a multi-hop query for pathways and biological processes related to cholesterol; clicking on nodes and edges shows their details on the right. When a result graph contains many intermediate nodes, the UI groups them based on type family, as can be seen for the 39 proteins (blue rectangle) connecting cholesterol to the cholesterol biosynthetic process node in **Figure 2**. Clicking on such a grouped node/edge lists all grouped items in the right panel, each of which is clickable for more details. Multi-hop queries are built using a graphical query builder, shown in **Figure 3**. Every query in the web interface is encoded in the page’s URL, so any query can be shared and rerun from its link; links for the queries behind the interface figures in this paper (Figures 1-4 and 8-10) are provided in Supplementary Data S6.

**Figure 2:**
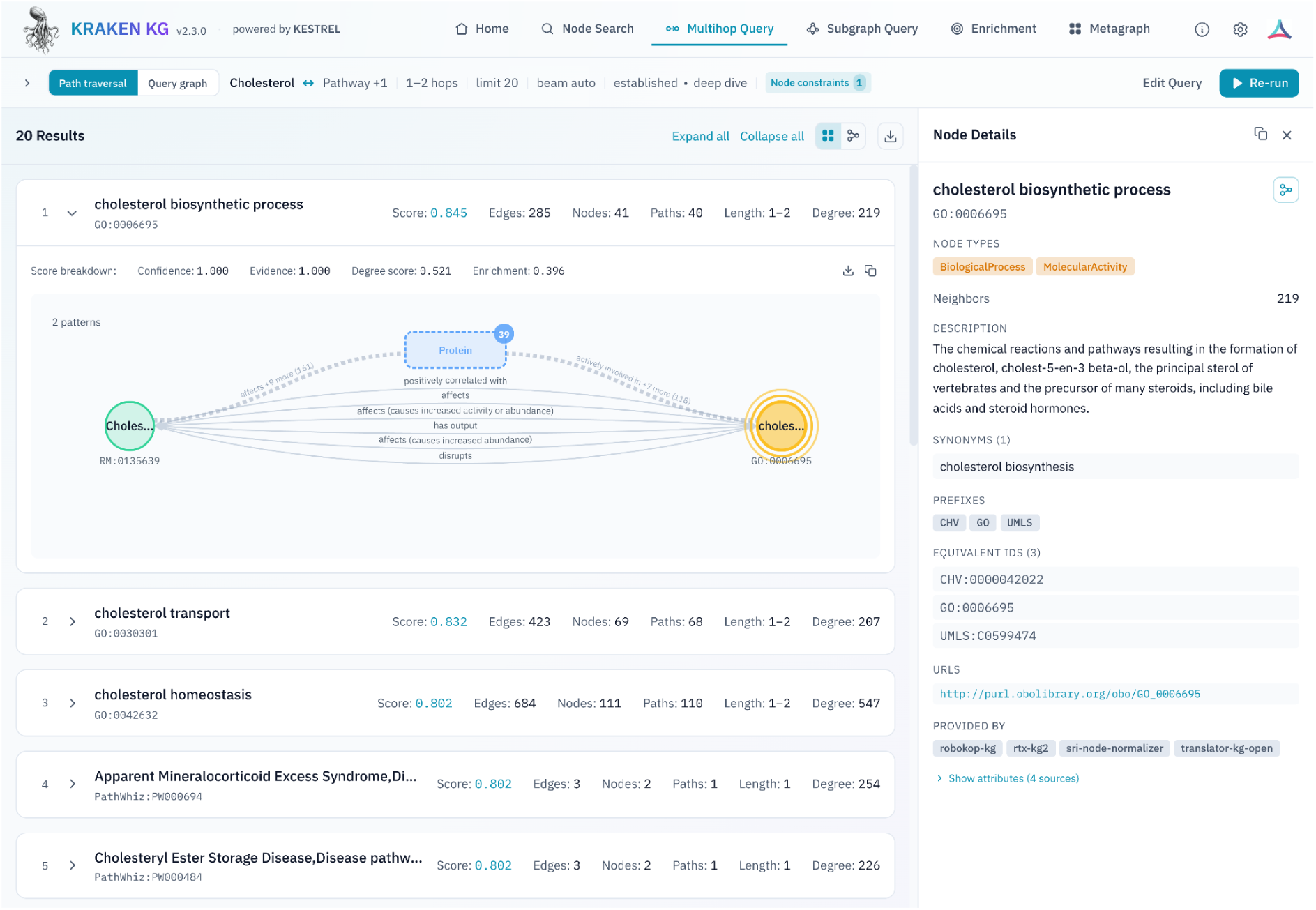
Example view of KRAKEN’s web interface showing results of a multi-hop query for pathways and biological processes connected to cholesterol; details for the first answer to this query (‘cholesterol biosynthetic process’) are displayed in the right side panel.

**Figure 3:**
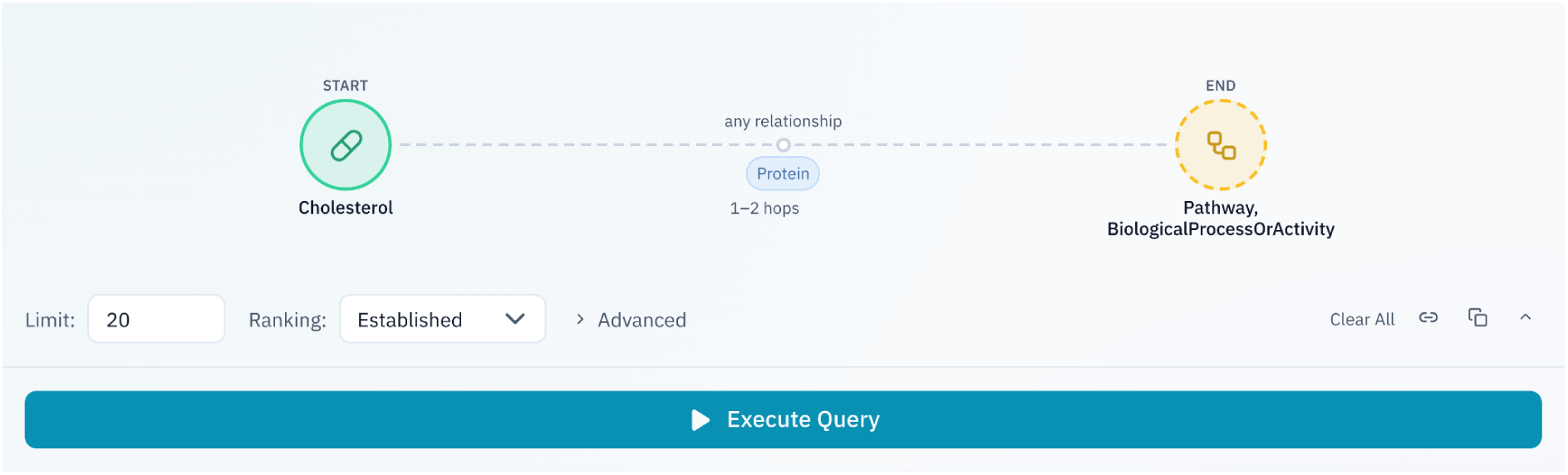
Example view of KRAKEN’s graphical query builder for multi-hop queries, showing the query submitted to produce the results shown in Figure 2.

**Figure 4:**
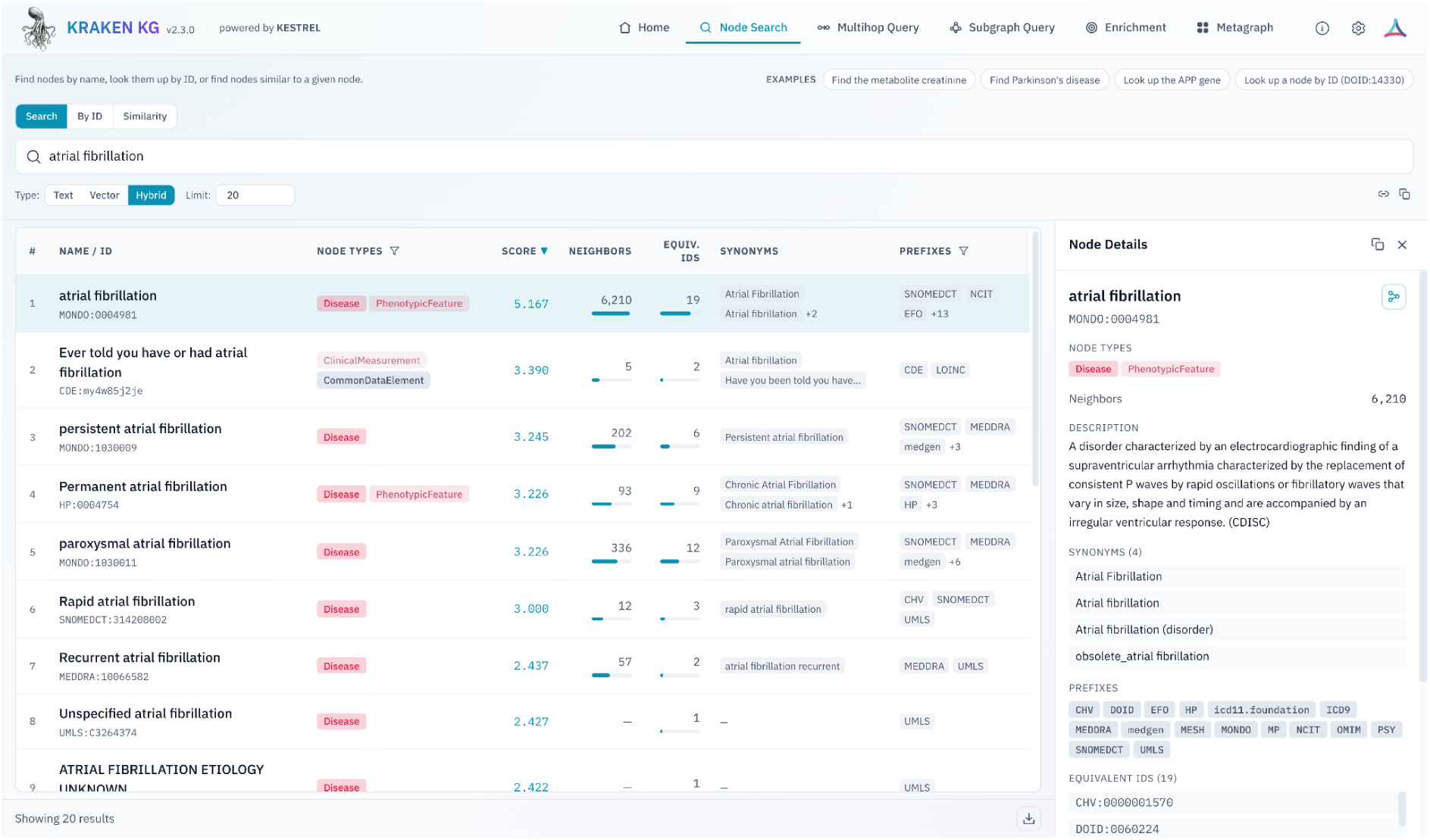
Example view of KRAKEN’s web interface showing a node search for ‘atrial fibrillation’. The first result is the canonical node for atrial fibrillation in KRAKEN; the second is a questionnaire data element from the NIH Common Data Elements repository.

**Figure 4** shows KRAKEN’s node search interface. It can be used standalone (as shown), and is also built into the query builders for multi-hop and subgraph queries, for easy selection of input nodes. It supports filtering by node type (hierarchically, per Biolink) and identifier vocabulary prefixes, and displays neighbor and equivalent ID counts for context.

### Analytical Tools

KRAKEN content is served by KESTREL, which provides various tools for querying KRAKEN, described below. Each analytical tool is exposed both through a REST endpoint and an MCP tool, with the web interface providing corresponding interactive access. These tools fall into two categories: node discovery tools, and graph analysis tools.

Node discovery tools, shown in **Table 2**, allow users to find entities of interest from free-text queries or similarity to known input nodes, using three search methods: lexical (text), semantic (vector), and hybrid (text + vector). The tools that rank by semantic similarity share a common set of node embeddings, generated (as of writing) with the all-MiniLM-L6-v2 sentence-transformer model from each node’s name and synonyms.

**Table 2.**
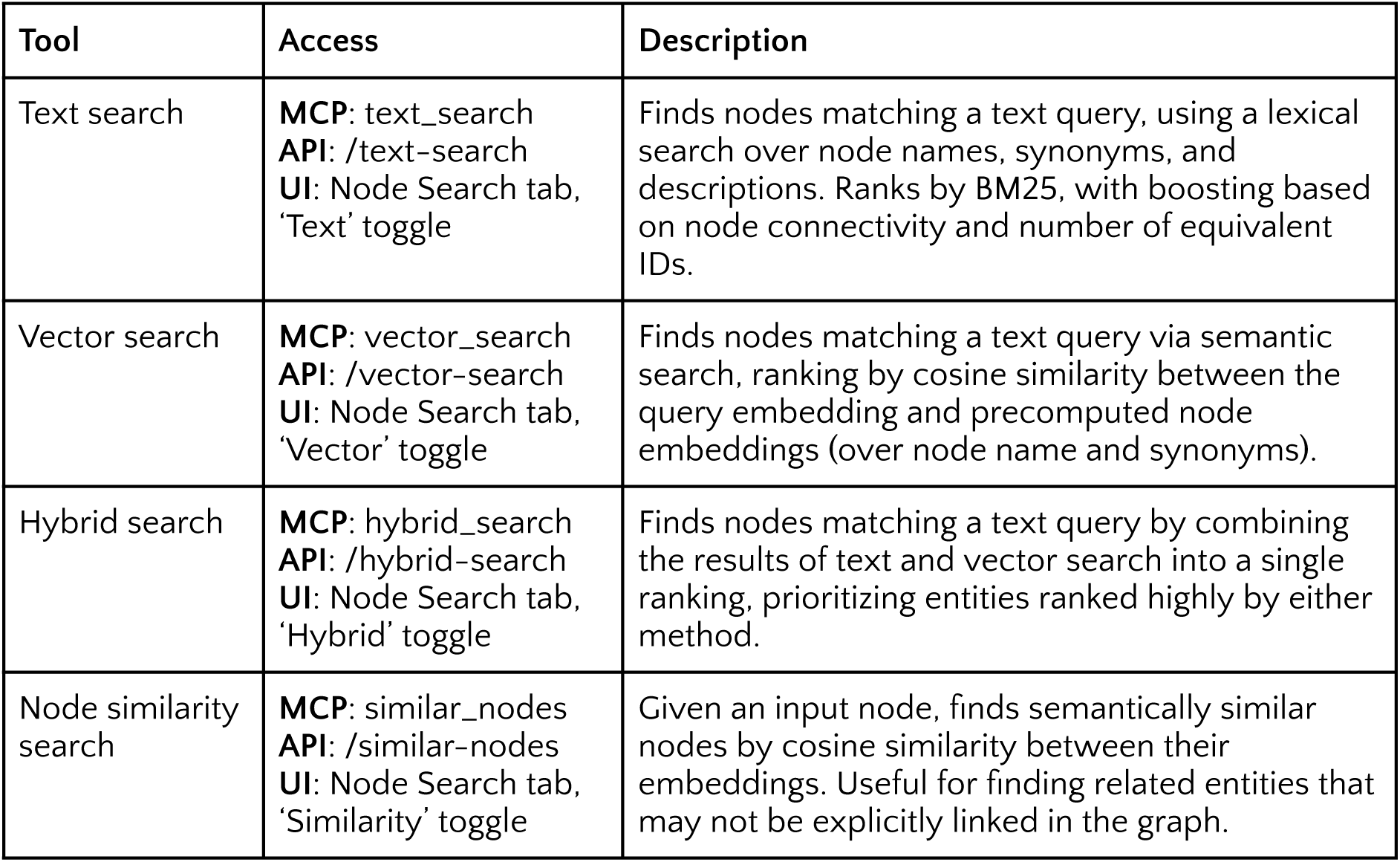
Node discovery tools available in KRAKEN. All support filtering by Biolink category (hierarchically) and identifier vocabulary prefix (e.g., HMDB).

Graph analysis tools, shown in **Table 3**, support exploration of relationships between entities and inference over their connections in the graph. Multi-hop reasoning, query graph matching, and subgraph extraction share a common constraint and ranking framework, described below. Enrichment analysis, which operates over the graph’s statistical structure rather than individual paths, has its own configuration described separately.

**Table 3.** Graph analysis tools available in KRAKEN. Multi-hop reasoning, query graph matching, and subgraph extraction share a common constraint and ranking framework (see main text).

| Tool | Access | Description |
| --- | --- | --- |
| Multi-hop reasoning | **MCP:** multi_hop_query<br>**API:** /multi-hop<br>**UI:** Multihop Query tab, 'Path traversal' view | Given one or more input entities, a target min/max path length, and optional constraints, returns paths that match the query pattern. Supports both singly-pinned queries (find connected entities from a start node) and doubly-pinned queries (find paths between start and end nodes). |
| Query graph matching | **MCP:** query_graph_query<br>**API:** /query-graph<br>**UI:** Multihop Query tab, 'Query graph' view | Matches a user-defined graph pattern against KRAKEN. Each query node can be pinned to specific entities or constrained by category, and each query edge matches a path of one or more hops. Pinned nodes can optionally be expanded to related concepts (e.g., subtypes, parent types, or equivalent terms). |
| Subgraph extraction | **MCP:** subgraph_query<br>**API:** /subgraph<br>**UI:** Subgraph Query tab | Given a set of input nodes, returns the connecting subgraph up to a specified depth. Ranks connecting nodes/paths and filters the subgraph to a user-specified size. |
| Enrichment analysis | **MCP:** kg_enrichment<br>**API:** /kg-enrichment<br>**UI:** Enrichment tab | Given a set of input nodes and optional target Biolink category and predicate, identifies target category nodes that are statistically enriched among the neighbors of the input set, using a degree-corrected permutation test. |

### Shared constraints and ranking (multi-hop, query graph, and subgraph tools)

These tools accept constraints on edge properties (predicate, knowledge level, agent type, knowledge source, qualifiers) and node properties (category, vocabulary prefix, connectivity/degree, chemical properties where applicable). All Biolink category and predicate constraints are hierarchical: constraining a category or predicate automatically includes its hyponyms. Results are ranked using a customizable model over four axes, with configurable preset weightings or user-specified weights. The four axes include 1) a confidence proxy derived from Biolink knowledge_level and agent_type properties on constituent edges; 2) an evidence proxy, based on the number of parallel edges between nodes; 3) a node degree component calculated as log-scaled degree, normalized within Biolink category; and 4) an enrichment component, favoring candidates connected to more of the input nodes than expected given their degree. Six presets are provided: Established (the default; favors well-supported, well-studied connections), Hidden Gems (high-quality evidence that has received little attention), Frontier (quality evidence connecting underexplored nodes), Deep Dive (well-supported connections specific to the query), Speculative (weaker evidence between well-known entities, for hypothesis generation), and Long Shot (weaker evidence between obscure entities); the weights each preset assigns are available at https://kestrel.krakenkg.com/api/traversal-options. These tools currently use beam search for graph traversal; the ranking model is applied both to intermediate traversal candidates and to final results, configurable separately for each.

### Enrichment analysis

This tool identifies nodes of a user-specified target category that are statistically over-represented among the neighbors of a set of input nodes. If a target category and predicate are not specified, they default to Biolink’s root NamedThing and related_to types, respectively (i.e., unrestricted); as with other tools, Biolink types are treated hierarchically. To avoid high-degree nodes dominating the enrichment results, we use a degree-corrected permutation test: for each candidate target, the observed number of input nodes connected by a matching edge is compared to a null distribution built from random, degree-matched node sets drawn from the background. By default, the background is the subset of KRAKEN nodes of the input category (e.g., Protein or SmallMolecule) with at least one edge matching the target predicate and category (e.g., participates_in, Pathway). Alternatively, the user may specify the background explicitly as a set of nodes (which are then restricted in the same way to those with a qualifying edge), to control for selection bias when the input nodes are drawn from a non-representative pool, such as a targeted assay panel. Empirical p-values are Benjamini-Hochberg corrected, and results include p-values, adjusted p-values, and fold enrichment.

## RESULTS

### Content and Scale

KRAKEN contains 26.5 million nodes and 120.7 million edges, spanning 62 Biolink node types and 113 edge types (v2.3.0). Full breakdowns of node and edge counts by Biolink type are provided in Supplementary **Tables S3** and **S4**, respectively. **Figure 5** visualizes KRAKEN’s meta-graph, showing rich cross-domain connectivity across genotypic, proteomic, chemical/metabolomic, functional, and phenotypic/clinical categories. Of KRAKEN’s 14,834 distinct meta-triples (e.g., SmallMolecule interacts_with Protein), 81% cross between different node type families.

**Figure 5:**
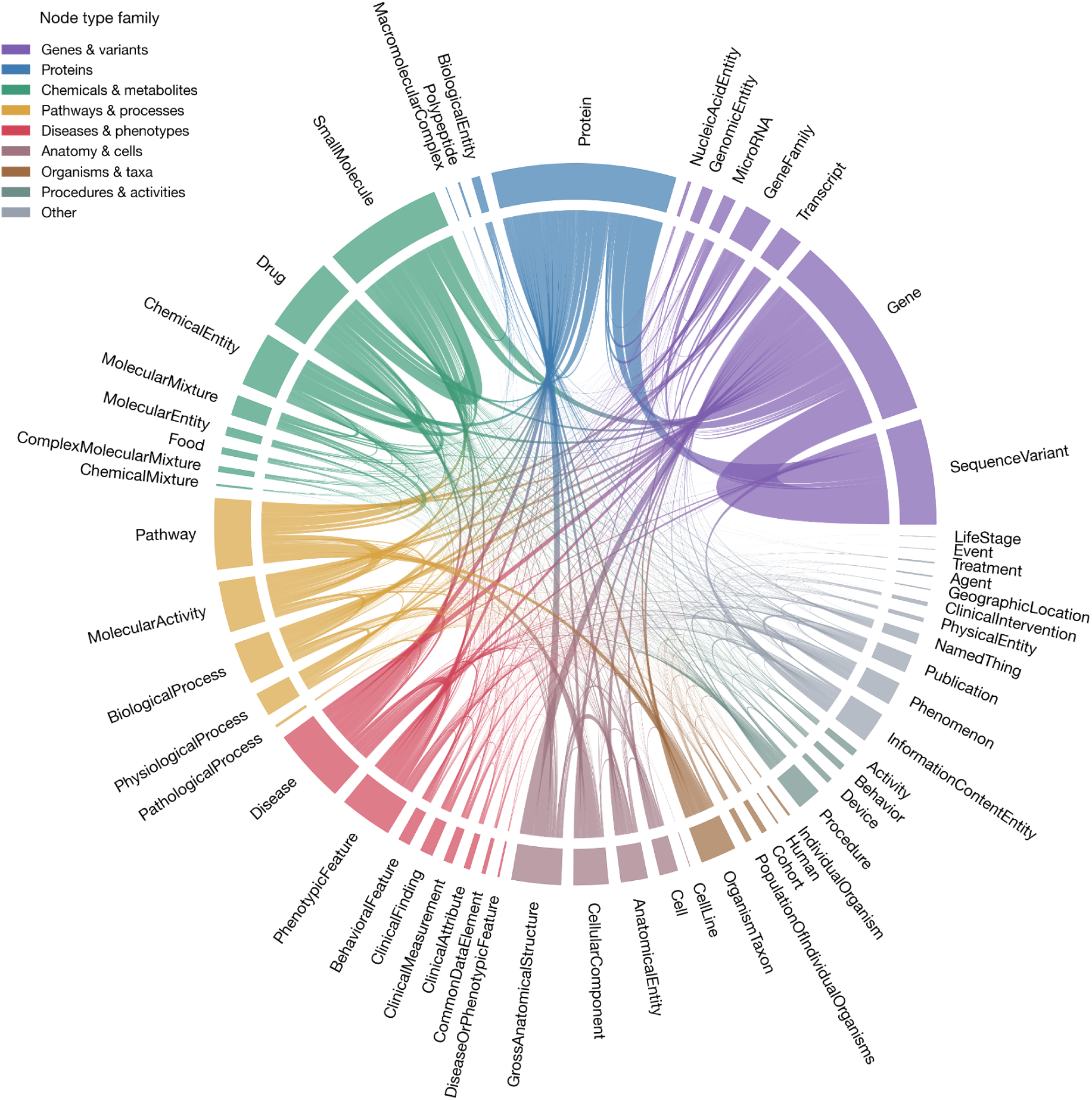
Chord diagram of KRAKEN’s meta-graph, showing schema-level connections between Biolink node types. Ribbon width represents the count of edges between each pair of node categories, square root-scaled to accommodate the range. For readability, only node category pairs with an edge count ≥100 are included in this visualization.

KRAKEN’s content is drawn from over 100 primary knowledge sources, including those ingested indirectly through other aggregator KGs. **Figure 6** highlights how these sources bridge to each other through shared entities in the integrated KRAKEN graph, enabling cross-source reasoning beyond that possible from any single source. The figure shows that sources tend to cluster more strongly within node type families, with certain sources such as Multiomics KG (Goetz et al., in preparation), HMDB (34), GO-Plus (35), DISEASES (36), and Uberon (37) emerging as stronger cross-domain bridges. Full lists of KRAKEN’s knowledge sources are provided in Supplementary **Tables S1** and **S2**.

**Figure 6:**
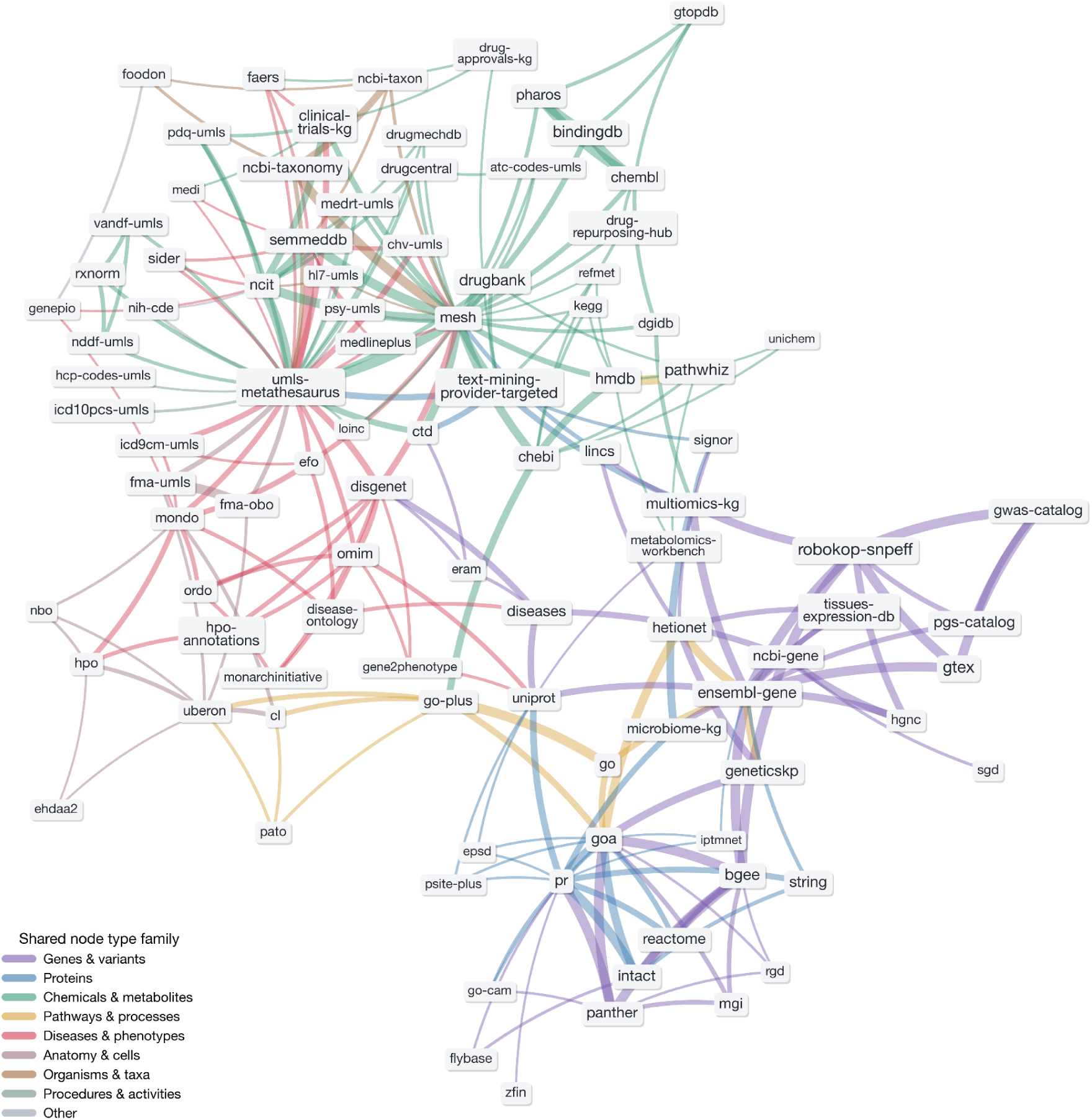
Force-directed network of primary knowledge source adjacency in KRAKEN, filtered for readability to show only the strongest connections. Each labeled pill is a primary knowledge source that contributes edges to KRAKEN; two sources are linked when edges they contribute meet at shared nodes. Without any restrictions this graph is very dense; here it is filtered to show only the top 3 connections per source that meet at ≥1,000 distinct nodes (sources without any such connections are excluded), yielding 240 links between 98 sources. Edge width is proportional to the number of distinct nodes the two sources share (linear, capped at the 90th percentile). Label size (log) scales with the total number of edges the source contributes to KRAKEN. Connections are colored per the dominant node type family of the shared nodes. Layout is by force-directed placement (Graphviz neato) with label de-overlap. Sources are referred to by their ‘infores’ ID (https://github.com/biolink/information-resource-registry) where available, with four exceptions made for clarity: multiomics-kg (corresponds to infores:multiomics-multiomics), microbiome-kg (infores:multiomics-microbiome), drug-approvals-kg (infores:multiomics-drugapprovals), and clinical-trials-kg (infores:multiomics-clinicaltrials). Aggregator sources (rtx-kg2, robokop-kg, translator-kg-open, sri-node-normalizer, ubergraph) are excluded.

The relationships provided by KRAKEN’s underlying sources were generated in varying ways, from manually-curated assertions to text-mined predictions; **Figure 7** visualizes the volume of edges according to Biolink knowledge level and agent type. The majority of KRAKEN’s edges come from either computational predictions (∼54 million) or manual curation/validation (∼31 million), with additional coverage from statistical associations, text mining, and logical entailment. KRAKEN’s analytical tools allow users to filter or weight results according to their tolerance for computational vs. curated evidence.

**Figure 7:**
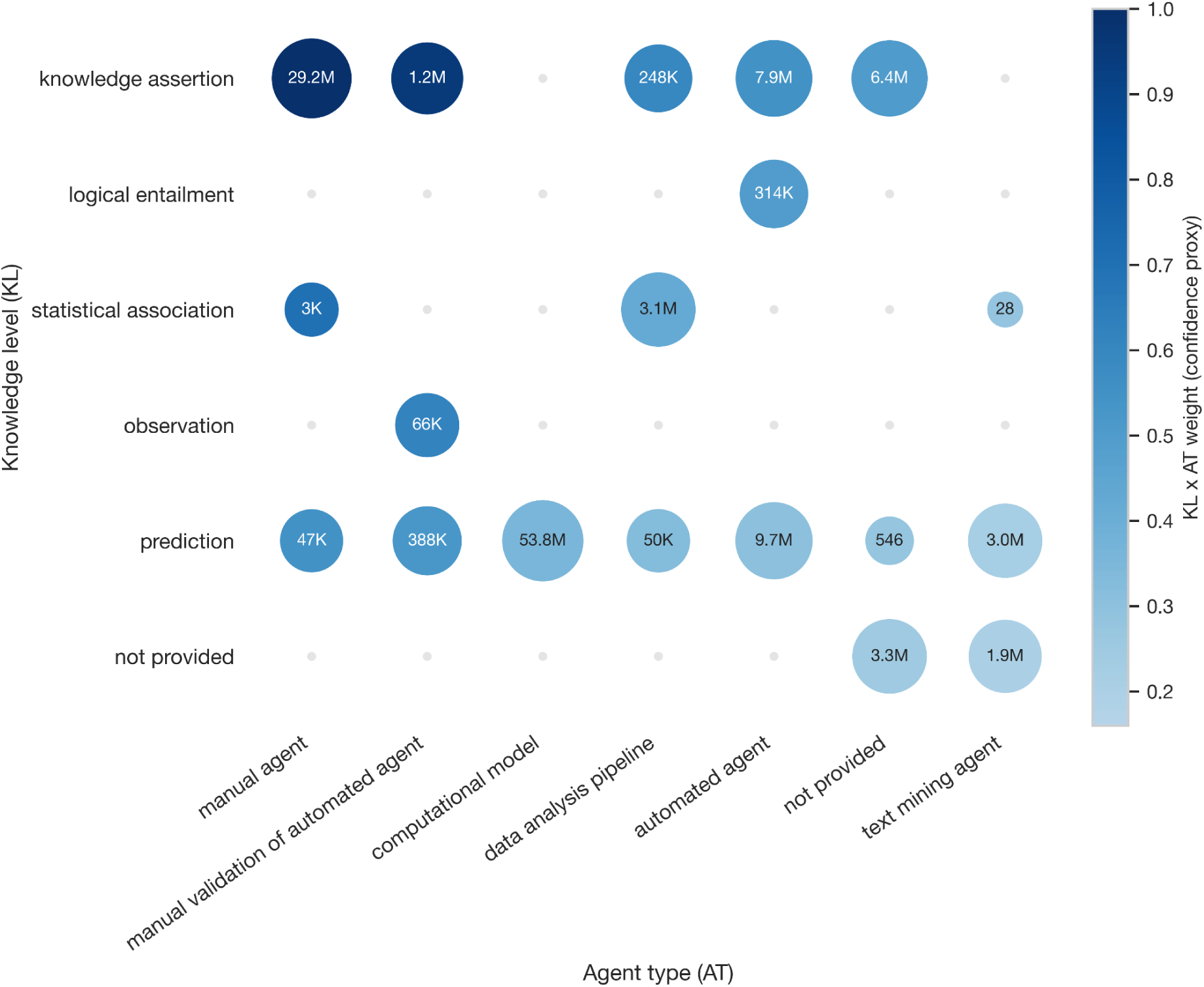
KRAKEN’s edges according to their Biolink knowledge_level and agent_type values. Dot size corresponds to edge count and is log-scaled. Dot color intensity corresponds to the confidence proxy used for such edges in query result ranking (described in Analytical Tools).

### Use Cases

#### Mechanistic interpretation of multiomic signature

A deep phenotyping analysis typically ends in a list of features that differ between groups or correlate with a phenotype, drawn from more than one assay and annotated in the vocabularies of the measurement platforms rather than in any ontology: metabolite names as reported by the mass spectrometry vendor, protein names by the affinity panel, clinical chemistries by local laboratory label. Two steps stand between that list and a mechanistic reading of it. The features must resolve to canonical entities, and the relationships among them must be retrieved and ranked. We illustrate both with 543 analytes associated with a combined measure of bBMI (13), comprising 356 metabolites, 149 proteins, and 38 clinical measures. KRAKEN’s node search resolved 482 labels with minimal manual curation, using text search with hybrid search as a fallback for synonyms text search alone did not recover. The 61 unresolved labels were almost entirely metabolites reported as unnamed spectral features, which carry no chemical name to match; the single named exception was the clinical laboratory OMEGA-3 INDEX, a composite without an unambiguous canonical entity. Enrichment analysis over the 88 more robustly selected signature proteins identified 400 pathways and biological processes at FDR below 0.05, including regulation of immune response, leukocyte migration, and chemokine-mediated signaling (**Figure 8**). The enriched terms converge on inflammatory signaling. Interpretation across omics is complicated by feature selection: sparse regression discards analytes that covary with retained ones, and targeted panels such as Olink’s Cardiovascular panel constrain what is measured at all. Graph relationships can restore the discarded connections. Subgraph extraction over the resolved node set, excluding disease, anatomical, information-content, and organism connectors, returned a connected structure of 660 nodes and 22,917 edges, comprising 245 genes and proteins, 201 drug-annotated chemicals, 171 chemicals without drug annotation, 35 processes, 5 clinical findings, and 3 cell types.

**Figure 8:**
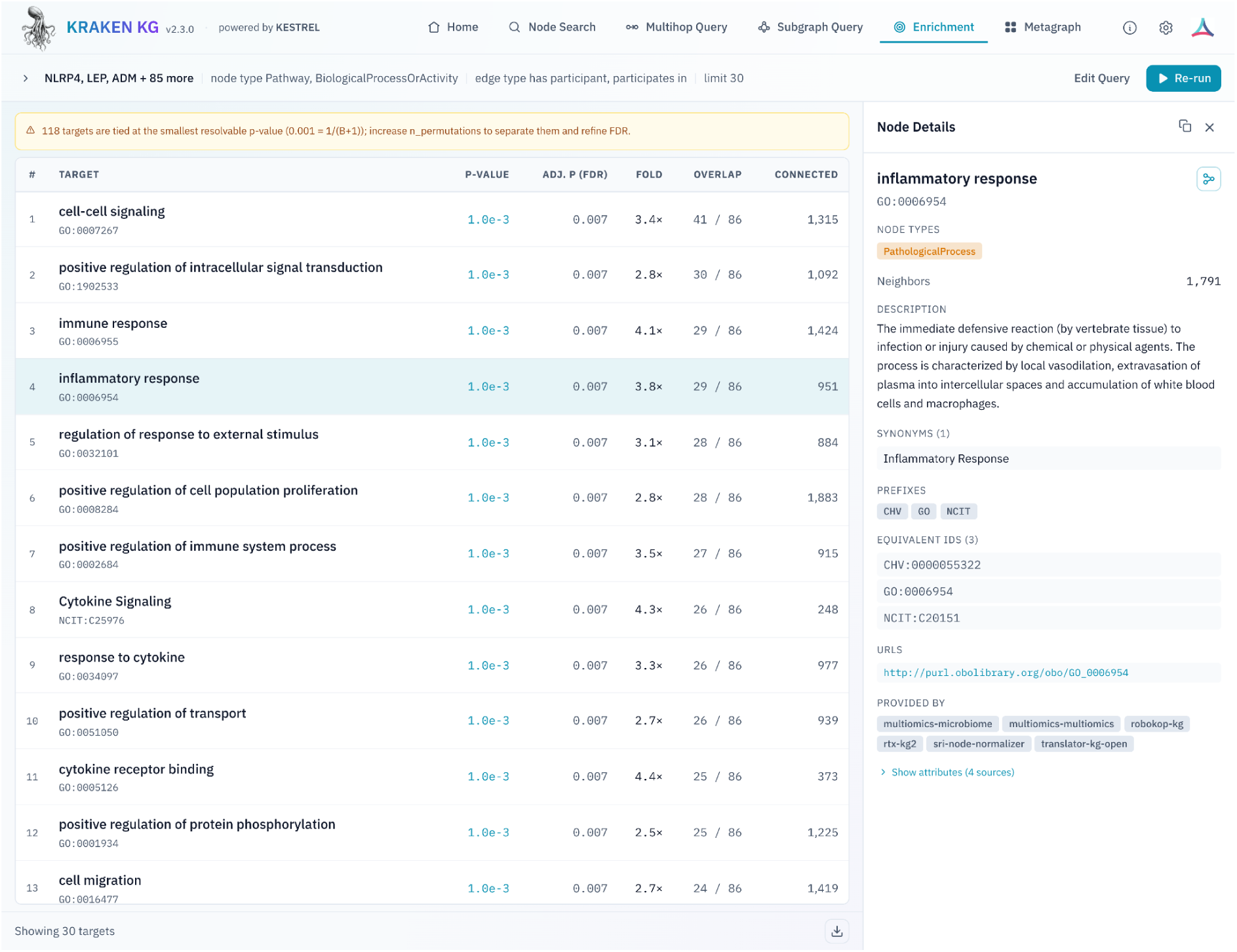
Pathways and biological processes enriched among signature proteins per KRAKEN’s KG enrichment tool.

The subgraph reorganized the signature along three axes. It extended the inflammatory component with IL6, TNF, IL1B, IL4, NFKB1, CXCL8, IFNG, IL10, STAT3, and TLR2/4 around an immune-response GO term hub, consolidating dispersed cytokines, chemokines, macrophage markers, and metabolic features into an adipose immunometabolic module (**Figure 9**); TNF signaling and pro-inflammatory adipose macrophages have established experimental and human links to obesity-associated insulin resistance (38, 39). It extended the endocrine component, whose largest model coefficients were LEP and FABP4, with INS, IGF1, IRS1, SOCS3, MAPK1/3, MTOR, GSK3B, PPARG, and ADIPOQ. It also added extracellular matrix and vascular remodeling analytes (TGFB1, TIMP1, ICAM1, SERPINE1/2, EDN1, AGT, PLG) connecting the measured MMPs, SELE, ADAMTS13, ADM, PLAT/PLAU, and renal signals, which implicate tissue remodeling, endothelial activation, and renin-angiotensin stress as secondary mechanisms. drug-annotated chemicals, 171 chemicals without drug Fourteen subgraph entities were analytes measured in the source study but discarded by the model in most training folds for covarying with a retained feature.

**Figure 9:**
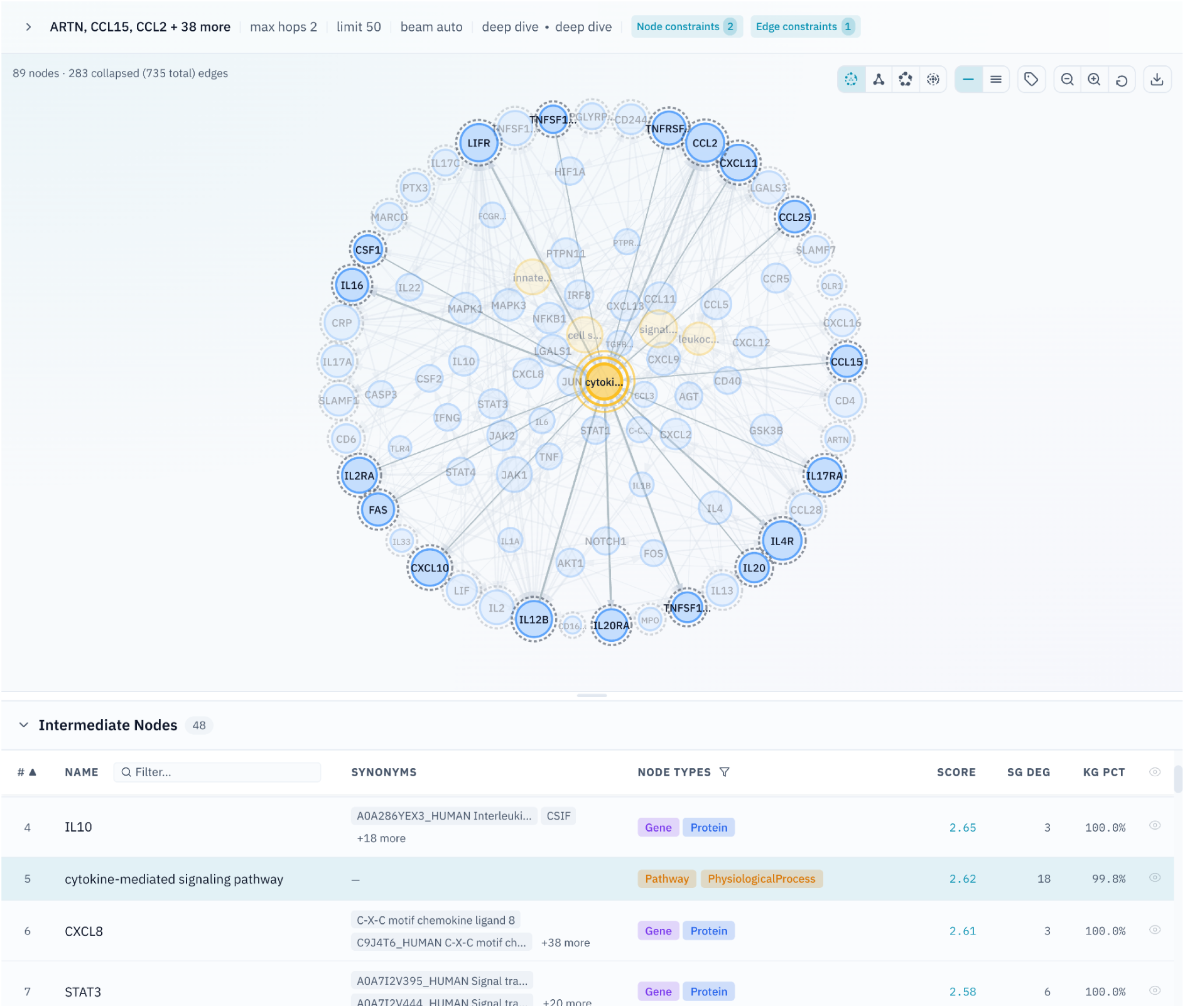
The inflammatory signaling module of the biological BMI subgraph in KRAKEN’s UI. A focused view of the immune and inflammatory neighborhood of the full use case subgraph, seeded from the measured inflammatory and immune proteins in the signature (outer ring of nodes) together with their shared protein and pathway connectors reachable within two steps over non-text-mined KRAKEN edges, limited to the top 48 connecting nodes.

### From polygenic risk to molecular intermediates and therapeutic hypotheses

A polygenic score quantifies inherited risk for a condition but doesn’t directly point to a mechanism or an intervention. Proteins measured in the same individuals whose circulating levels correlate with that score are candidate intermediates between the two, and a knowledge graph can propagate such a set forward to the compounds that act on those proteins and to whatever evidence exists that those compounds have been used clinically. The traversal is only possible in a graph where polygenic scores, proteins, compounds, diseases, and clinical evidence are all entities under one semantic layer. We have run this workflow using KRAKEN’s multi-hop functionality. Starting from plasma proteins whose levels correlated with polygenic risk for asthma in UK Biobank [2,923 proteins; (40)], a one-hop pattern query returned compounds connected to those proteins, with edges contributed by text-mined sources excluded so that any association recovered would not simply restate a published sentence. An optional second hop asked whether each compound had been approved for, used in, or trialled against asthma or a related condition. Three compounds are shown in **Figure 10**. Mepolizumab is an established asthma therapeutic and functions here as a positive control on the traversal. Marimastat, a matrix metalloproteinase inhibitor developed in oncology, had been evaluated in trials for conditions related to asthma; a subsequent literature search identified a randomized, double-blind, cross-over pilot study in twelve atopic asthmatic subjects in which marimastat reduced bronchial hyper-responsiveness to inhaled allergen while leaving exhaled nitric oxide, FEV1, symptom scores, and rescue inhaler use unchanged (41). The graph reached marimastat without access to that literature, since text-mined edges were excluded from the query, generating a potential repurposing candidate for further research. Marimastat nevertheless carries a text-mined edge asserting an asthma indication, so we repeated the search with an agent instructed to return only compounds whose supporting edges are text-mined at no step, executing KESTREL queries through the MCP server. Candidates were ranked by the specificity of the disease node bridging compound and indication, counted as the number of disease children that node carries, since a compound reaching its target only through a broad superclass has been connected to nothing in particular. Plasma CD30 (TNFRSF8) rose with asthma polygenic risk in the same cohort (β = 0.036, P = 2.4 × 10⁻⁷, Bonferroni-corrected P = 0.031, n = 20,341), and the one-hop query connected it to brentuximab vedotin (Adcetris), a CD30-directed antibody-drug conjugate used in Hodgkin lymphoma and anaplastic large-cell lymphoma (42). The compound reached asthma over the same lower respiratory tract bridge as marimastat, while no text-mined edge asserts an asthma indication for it. The CD30 axis is supported in asthma independently of the graph. A loss-of-function missense variant in TNFRSF8 protects against asthma by reducing CD30 on the cell surface and in soluble form (43), CD30-deficient mice are protected from TH2-mediated allergic airway inflammation (44, 45), and soluble CD30 is elevated in children with asthma (46, 47). Since TNFRSF8 is itself an asthma locus, the protein association may reflect cis effects of variants carried in the score rather than an independent intermediate. We present this as a target-level hypothesis rather than a treatment proposal. Brentuximab vedotin delivers a cytotoxic payload and carries pulmonary toxicity, so it is not itself a candidate in asthma. Instead, the traversal surfaced CD30 as a therapeutic axis from inherited risk alone.

**Figure 10:**
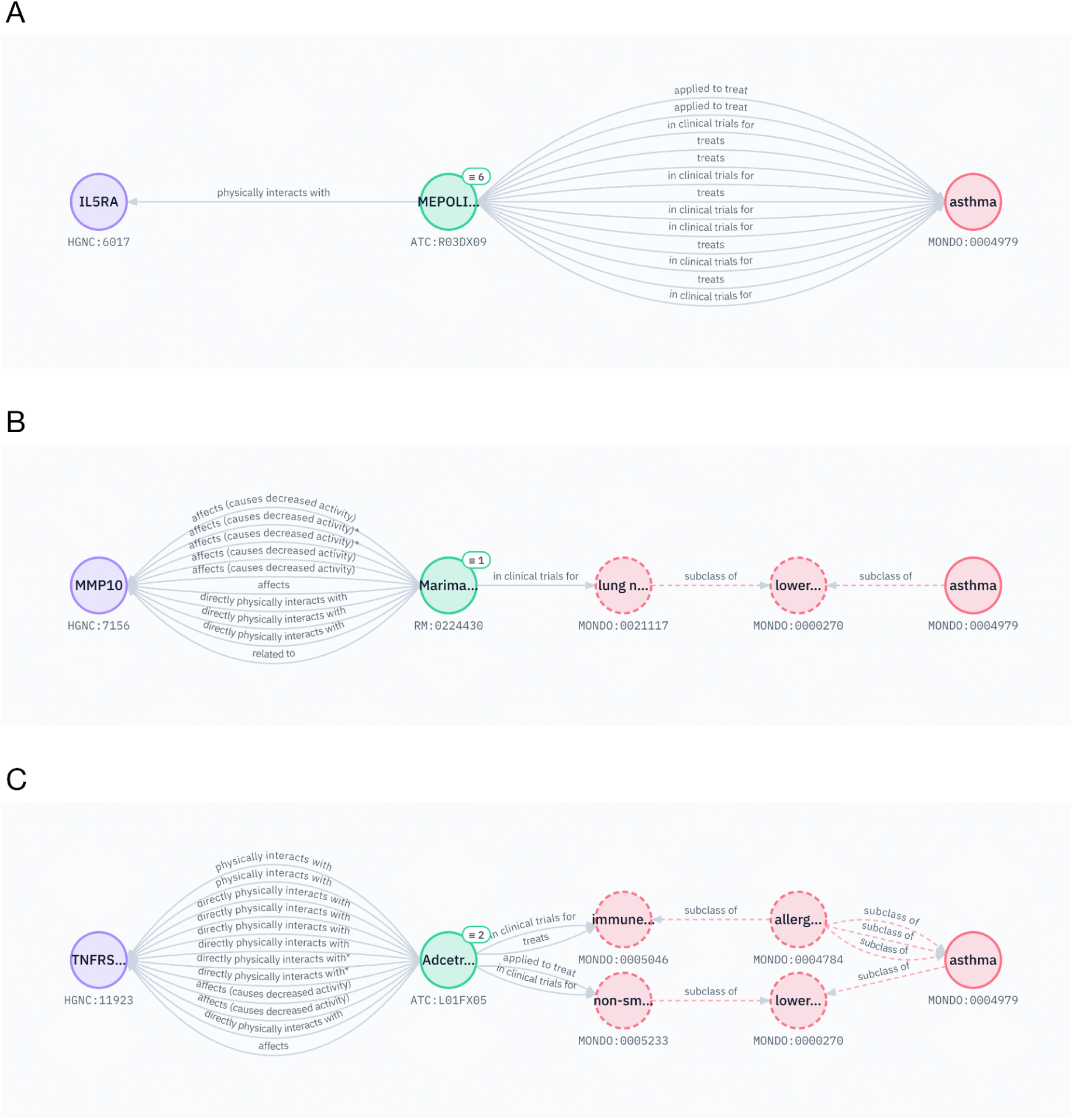
Three returned subgraphs connecting asthma PRS-correlated plasma proteins to compounds, with relevant clinical compound-to-disease edges included (retrieved using the query graph tool). Panel A shows the mepolizumab result, panel B the marimastat result, and panel C the brentuximab vedotin (Adcetris) result returned by the text-mining-free search. In all panels the compound node is shown together with its close-match nodes (≡ badge, with the number merged); in panels B and C asthma is expanded to related disease concepts (dashed nodes).

## DISCUSSION

KRAKEN was built to make multiomic and wellness data interpretable through a biomedical knowledge graph queryable by analysts and by analytical agents. The required content is scattered across resources assembled for other purposes, and much of it has never been represented as graph entities. Building it against the five properties set out above yields four capabilities: (1) analyte labels reported in the vocabularies of measurement platforms resolve against canonical entities, so harmonizing a feature list simplifies it to a query; (2) a cross-omic query can then begin at a metabolite or a lipid and reach e.g. proteins, pathways, and phenotypes without leaving one semantic layer, with metabolites carrying functional classifications alongside the structural ones on which most resources rely; (3) questions can start from entities that fill gaps in existing knowledge graphs, among them polygenic scores, standardized clinical data elements, laboratory observation codes, and derived measures such as BA and bBMI; and (4) an agent can execute these queries unattended and return claims a reader can trace to their sources, which is what distinguishes a grounded answer from a plausible one. Capabilities (1) and (3) depend on content absent from the comparator graphs, while (2) and (4) depend on that content sitting under a shared semantic layer with provenance preserved on each assertion. The analytical tools that support these queries are tailored to KRAKEN’s structure and reachable through all three interfaces: a web session, a programmatic call, and an agent.

KRAKEN consumes elements of the Translator ecosystem, including RTX-KG2, ROBOKOP, and Translator KG Open, and complements it with content oriented to multiomic and wellness analysis rather than disease mechanism and drug repurposing. The equivalences behind most canonical nodes derive from those assigned by the Translator SRI Node Normalizer, which KRAKEN draws on rather than deriving equivalences anew, supplying its own only where necessary. ORION, the pipeline constructing ROBOKOP (7), ingests and normalizes primary sources individually. KRAKEN instead builds largely on graphs that are already assembled and normalized. A full rebuild, entity resolution included, then runs on a laptop in under four hours with less than 48 GB of memory, and a schema change costs little to test. The build’s modularity lets a user include or exclude individual sources, and lets the build take Biolink-compliant community releases in place of the corresponding local ingests as they appear. Resources that operate at the level of the literature rather than the assertion are emerging in parallel. The NLM Linked Discoveries pilot assembles neighborhoods of related PubMed articles around a seed citation and surfaces citation context, retractions, and reviews (https://linkeddiscoveries.ncbi.nlm.nih.gov/). KRAKEN takes the complementary position of carrying evidence on the assertion itself, where knowledge_level and agent_type distinguish a curated claim from a text-mined one.

One design principle of KRAKEN is to preserve what its sources assert rather than forcing them to agree. Mapping a source predicate onto the nearest Biolink predicate discards the distinction the source drew, so KRAKEN keeps the original predicate and the source-specific properties on the edge. Where sources express one relation differently, some through complex predicates such as *upregulates* and others through qualified forms such as *affects* with direction and aspect, both are retained unreconciled. Merged identifiers likewise remain visible on the canonical node. We contend that this is the right default, since a user who disagrees with a hidden reconciliation cannot undo it, whereas a preserved distinction can be ignored. Metabolites illustrate the problem, since vocabularies draw the line between a compound class and its members inconsistently, and a user cannot tell from the graph which source a given merge relied on. KRAKEN also inherits the errors of its sources and validates no assertion, coverage across the Biolink hierarchy is uneven, with some categories drawn from a single source, and the wellness content is a starting point rather than exhaustive coverage of what a deep phenotyping study collects. Source acquisition remains semi-manual, since each addition needs a parser and a mapping decision. Node embeddings are computed from names and synonyms, so semantic search reflects nomenclature rather than function.

Work is under way on several of these limitations and additional features. Entity resolution is being rebuilt around clustering over identifiers, names, and structural descriptors, and in time a development of confidence probabilistic mapping scores, so that competing merges can coexist until evidence separates them. The two errors are not symmetric: a false merge contaminates every edge on both entities, whereas a missed merge only fragments the evidence, which argues for thresholds that favor precision on entities of high degree. Planned additions include further wellness measures, microbiome content, and standardized instruments, a topology-aware null model for enrichment, integrated issue reporting in the web user interface, and subgraph extraction that ranks candidate connecting structures rather than filtering them by size. The query and ranking methods underlying KESTREL will be reported separately. KRAKEN is in production use within the ARPA-H PATH program and will be maintained at its current URL, with quarterly releases, for at least five years.

## ACKNOWLEDGEMENTS

KRAKEN integrates content from many biomedical knowledge sources. We thank the developers and curators of these resources, including the NCATS Biomedical Data Translator Consortium and its member projects, whose knowledge graphs form key components of KRAKEN’s aggregated content. All sources are cited in **Table 1** and Supplementary **Tables S1** and **S2**. Sources with specific attribution requirements are listed below.

This work incorporates content from:

- The UMLS Metathesaurus produced by the U.S. National Library of Medicine. UMLS is available at https://www.nlm.nih.gov/research/umls/ under the UMLS Metathesaurus License Agreement.
- The NIH Common Data Elements Repository (https://cde.nlm.nih.gov/), National Library of Medicine, under the Open Data Commons Open Database License (ODbL).
- LOINC (http://loinc.org), copyright © Regenstrief Institute, Inc., available under the license at http://loinc.org/license. LOINC® is a registered trademark of Regenstrief Institute, Inc.

We thank John Earls for early inspiration that led to KRAKEN’s integrated MCP server.

We acknowledge use of generative AI (Anthropic’s Claude Code Opus 4.7, 4.8, and 5.5) to assist with code completion, comment generation, code debugging and manuscript preparation, specifically formatting and editing. All code, content and final wording have been reviewed and are solely the responsibility of the authors.

## AUTHOR CONTRIBUTIONS

Amy Glen: Conceptualization, Software, Methodology, Visualization, Writing—original draft, Writing—review & editing. Drew Witherington: Software. Trent Leslie: Methodology. Andrew Baumgartner: Methodology. Ashen Fernando: Methodology. Bhargav Vemuri: Methodology. Ornit Nahman: Methodology. Gwênlyn Glusman: Methodology, Writing—review & editing. Leroy Hood: Funding Acquisition. Lance Pflieger: Conceptualization, Methodology, Writing—original draft, Writing—review & editing. Noa Rappaport: Conceptualization, Methodology, Supervision, Writing—original draft, Writing—review & editing.

## SUPPLEMENTARY DATA

Supplementary Data S1-S6 are available with this preprint.

## CONFLICT OF INTEREST

None declared.

## FUNDING

This work was supported by an award from the Proactive Health Office of the Advanced Research Projects Agency for Health (ARPA-H) to the Personalized Analytics for Transforming Health (PATH) Project.

## DATA AVAILABILITY

KRAKEN is publicly available without registration or authentication at https://app.krakenkg.com and via the other interfaces described in Access Infrastructure. Content is provided under the licenses of the individual sources (see Table 1); no bulk data download is offered. Supplementary Table S1. provides a snapshot of primary knowledge source contributions as of KRAKEN v2.3.0. KRAKEN’s build code is publicly available under an MIT License on GitHub at https://github.com/Phenome-Health/kraken; the v2.3.0 release is archived on Zenodo at https://doi.org/10.5281/zenodo.23021807.

## Notes

### Competing Interest Statement

The authors have declared no competing interest.

### Summary of Updates

Updated from KRAKEN v2.1.0 to v2.3.0 (26.5 million nodes, 120.7 million edges). The web interface now supports shareable query links and downloads of results as Cytoscape-compatible JSON, tables, and images. Also adds five knowledge sources, a query graph matching tool, an enrichment component in ranking, expanded schema and backend descriptions, and a second candidate in the polygenic risk use case. All figures and statistics were regenerated, and the Supplementary Data were extended.

