## Supplementary Data S1-S6 for "KRAKEN: A provenance-tracked knowledge graph for multiomic and wellness research"

### KRAKEN knowledge graph: supplementary data

| | |
|---|---|
| Graph name | kraken |
| Graph version | 2.3.0 |
| Biolink Model version | 4.2.5 |
| Source metagraph | kraken_metagraph_2.3.0.json |
| Generated | 2026-10-07 |

#### Summary statistics

| | |
|---|---|
| Total nodes | 26,463,841 |
| Total edges | 120,705,263 |
| Unique node types (Biolink categories) | 62 |
| Unique vocabulary prefixes | 203 |
| Unique edge types (Biolink predicates) | 113 |
| Unique primary knowledge sources | 121 |
| Unique aggregator knowledge sources | 8 |
| Unique supporting data sources | 15 |

#### Contents


| | |
|---|---|
| S1. Primary knowledge sources | Every primary knowledge source and the number of edges attributed to it. |
| S2. Vocabulary prefixes | Every identifier prefix and the number of node identifiers that use it. |
| S3. Node types | Every node type and the number of nodes assigned to it. |
| S4. Edge types | Every edge type and the number of edges of that type. |
| S5. Example node and edge records | One node record and one edge record as they appear in KRAKEN's NDJSON files. |
| S6. Figure queries | Web interface queries used to generate the manuscript's figures. |


#### Notes

• In S2 (prefixes) and S3 (node types), the counts sum to MORE than the total number of nodes: a single node can carry multiple equivalent identifiers (hence multiple vocabulary prefixes) and multiple Biolink categories.

• In S1 (primary knowledge sources) and S4 (edge types), the counts sum to the total number of edges: each edge has exactly one primary knowledge source and one predicate.

• S1 includes four aggregator or identifier-mapping resources (robokop-kg, rtx-kg2, translator-kg-open and sri-node-normalizer): some edges are attributed directly to them rather than to an upstream source, for example close_match cross-references and, for robokop-kg, member_of edges.

### S1. Primary knowledge sources

| Primary knowledge source (infores CURIE) | Edge count |
|---|---|
| infores:robokop-snpeff | 35,268,451 |
| infores:gtex | 18,540,748 |
| infores:ubergraph | 10,031,065 |
| infores:pathwhiz | 8,964,361 |
| infores:robokop-kg | 5,424,755 |
| infores:bgee | 5,232,785 |
| infores:drugbank | 2,975,291 |
| infores:bindingdb | 2,921,481 |
| infores:hmdb | 2,382,640 |
| infores:ensembl-gene | 2,347,468 |
| infores:semmeddb | 1,857,400 |
| infores:intact | 1,849,004 |
| pgs-catalog | 1,507,105 |
| infores:goa | 1,383,685 |
| infores:text-mining-provider-targeted | 1,370,709 |
| infores:diseases | 1,175,441 |
| infores:string | 1,103,152 |
| infores:ncit | 994,224 |
| infores:hetionet | 910,229 |
| infores:geneticskp | 891,056 |
| infores:multiomics-clinicaltrials | 833,006 |
| infores:mesh | 756,665 |
| infores:panther | 744,461 |
| infores:ncbi-taxonomy | 742,788 |
| infores:reactome | 693,105 |
| infores:pr | 627,504 |
| infores:umls-metathesaurus | 600,480 |
| infores:lincs | 544,970 |
| infores:multiomics-multiomics | 480,514 |
| infores:gwas-catalog | 445,029 |
| infores:disgenet | 407,518 |
| infores:rtx-kg2 | 401,458 |
| infores:pharos | 399,980 |
| infores:chebi | 363,603 |
| infores:chembl | 348,513 |
| infores:hpo-annotations | 311,921 |
| infores:ncbi-gene | 296,110 |
| infores:omim | 273,375 |
| infores:sider | 271,625 |
| infores:faers | 268,383 |
| infores:sri-node-normalizer | 258,574 |
| infores:fma-umls | 232,846 |
| infores:mgi | 226,531 |
| infores:go-plus | 222,058 |

### S1. Primary knowledge sources  (continued)

| Primary knowledge source (infores CURIE) | Edge count |
|---|---|
| infores:ctd | 199,952 |
| infores:icd10pcs-umls | 191,463 |
| translator-kg-open | 178,591 |
| infores:go | 157,759 |
| infores:tissues-expression-db | 157,013 |
| infores:rxnorm | 149,757 |
| infores:fma-obo | 119,466 |
| infores:medrt-umls | 114,755 |
| infores:efo | 99,473 |
| infores:mondo | 89,241 |
| infores:drugcentral | 88,812 |
| infores:multiomics-microbiome | 88,231 |
| infores:signor | 85,072 |
| infores:uniprot | 78,108 |
| infores:uberon | 64,994 |
| infores:hpo | 63,170 |
| infores:vandf-umls | 59,761 |
| infores:dgidb | 58,471 |
| infores:foodon | 54,429 |
| infores:ordo | 52,896 |
| infores:gtopdb | 52,565 |
| infores:zfin | 45,653 |
| infores:cl | 43,543 |
| infores:nddf-umls | 34,168 |
| infores:hgnc | 31,797 |
| infores:psy-umls | 31,677 |
| infores:icd9cm-umls | 31,106 |
| infores:ncbi-taxon | 31,069 |
| infores:pdq-umls | 26,256 |
| nih-cde | 22,686 |
| infores:chv-umls | 22,555 |
| infores:flybase | 22,301 |
| infores:disease-ontology | 21,923 |
| infores:pato | 19,454 |
| infores:drug-repurposing-hub | 19,388 |
| infores:icees-kg | 13,945 |
| infores:medlineplus | 13,522 |
| infores:kegg | 13,323 |
| infores:drugmechdb | 13,107 |
| infores:ehdaa2 | 13,080 |
| infores:eram | 12,938 |
| infores:rgd | 12,829 |
| infores:sgd | 12,120 |
| infores:nbo | 11,244 |

### S1. Primary knowledge sources (continued)

| Primary knowledge source (infores CURIE) | Edge count |
|---|---|
| infores:genepio | 10,386 |
| infores:hl7-umls | 9,614 |
| infores:monarchinitiative | 8,676 |
| infores:hcp-codes-umls | 8,215 |
| infores:atc-codes-umls | 7,058 |
| infores:psite-plus | 6,965 |
| infores:refmet | 5,898 |
| infores:metabolomics_workbench | 5,503 |
| infores:epsd | 5,370 |
| infores:multiomics-drugapprovals | 5,174 |
| infores:go-cam | 4,252 |
| infores:medi | 3,911 |
| infores:gene2phenotype | 3,046 |
| infores:loinc | 2,225 |
| infores:ccidb | 2,222 |
| infores:iptmnet | 1,893 |
| infores:wombat-pk | 1,769 |
| infores:mi | 1,640 |
| infores:unichem | 1,614 |
| lipidmaps | 1,461 |
| infores:pdsp | 734 |
| infores:dda | 352 |
| infores:ino | 343 |
| infores:bspo | 342 |
| infores:ro | 195 |
| infores:amyco | 181 |
| isb-long-covid | 180 |
| infores:drugmatrix | 99 |
| infores:cohd | 84 |
| infores:pdsp-ki | 59 |
| infores:bfo | 34 |
| biological-age | 22 |
| biological-bmi | 16 |

### S2. Vocabulary prefixes

| Vocabulary prefix | Node count |
|---|---|
| DBSNP | 5,521,791 |
| CAID | 5,102,613 |
| HGVS | 4,839,615 |
| ENSEMBL | 3,768,512 |
| INCHIKEY | 3,660,050 |
| PUBCHEM.COMPOUND | 3,544,169 |
| UMLS | 3,375,381 |
| UniProtKB | 3,337,627 |
| CHEMBL.COMPOUND | 2,927,119 |
| NCBITaxon | 2,873,168 |
| NCBIGene | 2,404,368 |
| MGI | 640,462 |
| CAS | 491,052 |
| MESH | 359,089 |
| PR | 255,575 |
| CHEBI | 222,377 |
| LOINC | 218,526 |
| HMDB | 218,198 |
| RM | 206,981 |
| SMILES | 195,864 |
| ICD10PCS | 191,277 |
| NCIT | 182,722 |
| REACT | 179,966 |
| PathWhiz.Reaction | 175,039 |
| UNII | 167,977 |
| RGD | 151,160 |
| RXCUI | 123,027 |
| SNOMEDCT | 121,136 |
| VGNC | 112,849 |
| SMPDB | 110,229 |
| PathWhiz | 110,217 |
| RXNORM | 107,563 |
| OMIM | 106,230 |
| FMA | 105,021 |
| CLINVARVARIANT | 81,182 |
| PathWhiz.Compound | 78,599 |
| RHEA | 74,232 |
| MEDDRA | 60,471 |
| CHV | 57,790 |
| LM | 56,626 |
| WB | 49,533 |
| Xenbase | 49,022 |
| HGNC | 45,194 |
| GO | 44,733 |

### S2. Vocabulary prefixes (continued)

| Vocabulary prefix | Node count |
|---|---|
| CLO | 38,897 |
| AraPort | 38,127 |
| ZFIN | 38,023 |
| FB | 32,179 |
| NDDF | 31,391 |
| VANDF | 30,353 |
| MONDO | 28,070 |
| FOODON | 26,634 |
| PANTHER.FAMILY | 26,140 |
| CGNC | 26,053 |
| ICD9 | 22,405 |
| CDE | 22,308 |
| medgen | 21,452 |
| ComplexPortal | 20,579 |
| KEGG.COMPOUND | 19,735 |
| HP | 19,518 |
| DRUGBANK | 19,420 |
| EFO | 16,238 |
| CHEMBL.TARGET | 15,394 |
| UBERON | 14,973 |
| orphanet | 14,948 |
| MP | 14,751 |
| BGD | 14,434 |
| mirbase | 13,998 |
| dictyBase | 13,892 |
| GTOPDB | 13,863 |
| PDQ | 13,325 |
| KEGG.DRUG | 12,618 |
| SLM | 12,557 |
| PomBase | 12,420 |
| DOID | 12,294 |
| KEGG.REACTION | 12,242 |
| KEGG.GLYCAN | 11,232 |
| OBA | 11,168 |
| PathWhiz.ProteinComplex | 8,882 |
| KEGG.ENZYME | 8,235 |
| EMAPA | 8,098 |
| SGD | 8,097 |
| PSY | 7,959 |
| KEGG | 7,471 |
| EC | 7,375 |
| HCPCS | 7,327 |
| ttd.target | 7,147 |
| ATC | 6,564 |

#### S2. Vocabulary prefixes (continued)

| Vocabulary prefix | Node count |
|---|---|
| DrugCentral | 4,996 |
| dictybase.gene | 4,555 |
| OBO | 4,207 |
| icd11.foundation | 4,058 |
| ecogene | 4,042 |
| CL | 3,346 |
| LIPIDBANK | 2,747 |
| ICD10 | 2,586 |
| EHDAA2 | 2,459 |
| MA | 2,450 |
| HGNC.FAMILY | 2,323 |
| ZFA | 2,076 |
| miRBase | 1,915 |
| CHEMBL.MECHANISM | 1,887 |
| PATO | 1,881 |
| MI | 1,631 |
| GENEPIO | 1,596 |
| CDNO | 1,100 |
| NBO | 932 |
| ENVO | 710 |
| OMIM.PS | 594 |
| PLANTFA | 556 |
| PO | 488 |
| FBbt | 459 |
| ECTO | 430 |
| XAO | 427 |
| PathWhiz.Bound | 416 |
| PGS | 412 |
| PathWhiz.ElementCollection | 340 |
| EnsemblGenomes | 280 |
| PathWhiz.NucleicAcid | 251 |
| OBI | 237 |
| SO | 208 |
| UO | 191 |
| PANTHER.PATHWAY | 175 |
| MOD | 170 |
| BSPO | 140 |
| AEO | 135 |
| DDANAT | 131 |
| MAXO | 115 |
| FAO | 112 |
| BTO | 89 |
| MPATH | 76 |
| STATO | 71 |

### S2. Vocabulary prefixes (continued)

| Vocabulary prefix | Node count |
|---|---|
| OMIT | 63 |
| ONS | 63 |
| ARO | 55 |
| IAO | 54 |
| WBbt | 48 |
| CARO | 46 |
| EO | 43 |
| ECOCORE | 41 |
| KEGG.DISEASE | 40 |
| IDO | 39 |
| SEMMEDDB | 38 |
| BFO | 35 |
| TCDB | 23 |
| umls_source | 23 |
| SIO | 16 |
| MF | 14 |
| PCO | 14 |
| OMRSE | 13 |
| OGG | 13 |
| OPL | 13 |
| MFOMD | 12 |
| identifiers_org_registry | 12 |
| COB | 11 |
| OGMS | 10 |
| dbpedia | 10 |
| CHMO | 8 |
| TRANS | 7 |
| PW | 6 |
| AGRO | 6 |
| icd11 | 6 |
| RO | 5 |
| FLOPO | 4 |
| BAO | 3 |
| MMO | 3 |
| SYMP | 3 |
| VCARD | 2 |
| CEPH | 2 |
| ExO | 2 |
| EDAM-DATA | 2 |
| RTX | 2 |
| TO | 2 |
| VT | 2 |
| XCO | 1 |
| ZEA | 1 |

| Vocabulary prefix | Node count |
|---|---|
| UPHENO | 1 |
| HANCESTRO | 1 |
| OMIABIS | 1 |
| GAZ | 1 |
| ICO | 1 |
| FLU | 1 |
| BIOAGE | 1 |
| BIOBMI | 1 |
| CVDO | 1 |
| ClinicalTrialsKG | 1 |
| DGIdb | 1 |
| DisGeNET | 1 |
| DrugApprovalsKG | 1 |
| EPO | 1 |
| ERO | 1 |
| EUPATH | 1 |
| FBdv | 1 |
| FIX | 1 |
| GEO | 1 |
| JensenLab | 1 |
| KEGG_source | 1 |
| NCRO | 1 |
| OMP | 1 |
| TypOn | 1 |
| UNICHEM_source | 1 |
| WBls | 1 |
| biolink_download_source | 1 |

### S3. Node types

| Node type (Biolink category) | Node count |
| --- | --- |
| biolink:SequenceVariant | 10,408,391 |
| biolink:SmallMolecule | 3,774,924 |
| biolink:Gene | 3,300,131 |
| biolink:Protein | 3,287,416 |
| biolink:OrganismTaxon | 2,946,876 |
| biolink:Procedure | 452,717 |
| biolink:Disease | 363,683 |
| biolink:MolecularActivity | 350,017 |
| biolink:Phenomenon | 333,161 |
| biolink:ChemicalEntity | 326,861 |
| biolink:Drug | 316,955 |
| biolink:Transcript | 269,901 |
| biolink:AnatomicalEntity | 149,579 |
| biolink:MolecularMixture | 141,405 |
| biolink:PhenotypicFeature | 127,848 |
| biolink:InformationContentEntity | 119,308 |
| biolink:Pathway | 119,198 |
| biolink:ClinicalAttribute | 111,448 |
| biolink:ClinicalFinding | 104,489 |
| biolink:NamedThing | 82,664 |
| biolink:Device | 79,633 |
| biolink:ClinicalMeasurement | 76,002 |
| biolink:Publication | 54,536 |
| biolink:BiologicalProcess | 43,635 |
| biolink:GenomicEntity | 42,568 |
| biolink:BiologicalEntity | 40,955 |
| biolink:CellLine | 39,914 |
| biolink:ClinicalIntervention | 36,758 |
| biolink:GeneFamily | 25,834 |
| biolink:CommonDataElement | 21,658 |
| biolink:MacromolecularComplex | 19,246 |
| biolink:GrossAnatomicalStructure | 14,311 |
| biolink:MolecularEntity | 13,993 |
| biolink:PopulationOfIndividualOrganisms | 13,635 |
| biolink:PhysicalEntity | 13,171 |
| biolink:Activity | 12,761 |
| biolink:Food | 9,740 |
| biolink:Cell | 9,423 |
| biolink:CellularComponent | 8,771 |
| biolink:PathologicalProcess | 8,131 |
| biolink:DiseaseOrPhenotypicFeature | 6,732 |
| biolink:Behavior | 4,888 |
| biolink:GeographicLocation | 4,868 |
| biolink:Agent | 4,686 |

### S3. Node types (continued)

| Node type (Biolink category) | Node count |
|---|---|
| biolink:PhysiologicalProcess | 4,345 |
| biolink:Cohort | 2,555 |
| biolink:MicroRNA | 1,917 |
| biolink:NucleicAcidEntity | 981 |
| biolink:Polypeptide | 960 |
| biolink:ComplexMolecularMixture | 584 |
| biolink:IndividualOrganism | 523 |
| biolink:ChemicalMixture | 456 |
| biolink:BehavioralFeature | 320 |
| biolink:Event | 298 |
| biolink:Treatment | 74 |
| biolink:RetrievalSource | 70 |
| biolink:EnvironmentalProcess | 67 |
| biolink:Human | 46 |
| biolink:RNAProduct | 44 |
| biolink:NoncodingRNAProduct | 36 |
| biolink:MaterialSample | 13 |
| biolink:LifeStage | 10 |

### S4. Edge types

| Edge type (Biolink predicate) | Edge count |
|---|---|
| biolink:is_nearby_variant_of | 31,403,914 |
| biolink:affects | 22,774,079 |
| biolink:subclass_of | 14,057,783 |
| biolink:has_participant | 6,275,180 |
| biolink:expressed_in | 5,514,071 |
| biolink:member_of | 5,188,257 |
| biolink:occurs_in | 4,450,168 |
| biolink:is_non_coding_variant_of | 3,789,454 |
| biolink:related_to | 3,344,867 |
| biolink:interacts_with | 3,092,226 |
| biolink:directly_physically_interacts_with | 2,305,712 |
| biolink:gene_associated_with_condition | 1,738,902 |
| biolink:physically_interacts_with | 1,667,251 |
| biolink:close_match | 1,539,137 |
| biolink:contributes_to | 1,538,574 |
| biolink:actively_involved_in | 1,227,058 |
| biolink:located_in | 916,551 |
| biolink:has_phenotype | 824,506 |
| biolink:regulates | 821,502 |
| biolink:in_clinical_trials_for | 786,044 |
| biolink:has_part | 766,976 |
| biolink:has_input | 737,035 |
| biolink:has_output | 548,502 |
| biolink:treats_or_applied_or_studied_to_treat | 467,616 |
| biolink:genetic_association | 415,028 |
| biolink:in_taxon | 362,344 |
| biolink:causes | 291,166 |
| biolink:catalyzes | 274,921 |
| biolink:transcribed_from | 269,941 |
| biolink:enables | 243,647 |
| biolink:coexists_with | 243,580 |
| biolink:condition_associated_with_gene | 238,014 |
| biolink:associated_with | 216,353 |
| biolink:has_adverse_event | 202,110 |
| biolink:correlated_with | 175,611 |
| biolink:genetically_interacts_with | 147,139 |
| biolink:genetically_associated_with | 145,818 |
| biolink:has_side_effect | 139,765 |
| biolink:manifestation_of | 131,266 |
| biolink:associated_with_resistance_to | 128,642 |
| biolink:acts_upstream_of_or_within | 93,594 |
| biolink:precedes | 86,851 |
| biolink:active_in | 82,961 |
| biolink:orthologous_to | 76,483 |

### S4. Edge types (continued)

| Edge type (Biolink predicate) | Edge count |
|---|---|
| biolink:applied_to_treat | 76,339 |
| biolink:associated_with_sensitivity_to | 73,474 |
| biolink:treats | 67,556 |
| biolink:colocalizes_with | 60,050 |
| biolink:biomarker_for | 52,182 |
| biolink:translates_to | 48,368 |
| biolink:chemically_similar_to | 46,163 |
| biolink:overlaps | 38,126 |
| biolink:gene_product_of | 37,153 |
| biolink:disrupts | 36,323 |
| biolink:is_sequence_variant_of | 36,313 |
| biolink:part_of | 35,041 |
| biolink:is_missense_variant_of | 31,058 |
| biolink:contraindicated_in | 31,045 |
| biolink:diagnoses | 29,253 |
| biolink:derives_from | 28,698 |
| biolink:is_synonymous_variant_of | 28,056 |
| biolink:predisposes_to_condition | 24,213 |
| biolink:produces | 22,534 |
| biolink:has_active_ingredient | 17,884 |
| biolink:preventative_for_condition | 17,758 |
| biolink:is_splice_site_variant_of | 13,473 |
| biolink:has_member | 11,570 |
| biolink:coexpressed_with | 11,390 |
| biolink:lacks_part | 8,344 |
| biolink:capable_of | 7,774 |
| biolink:develops_from | 7,540 |
| biolink:similar_to | 6,934 |
| biolink:positively_correlated_with | 6,895 |
| biolink:target_for | 6,216 |
| biolink:disease_has_location | 6,145 |
| biolink:affects_response_to | 5,443 |
| biolink:acts_upstream_of | 4,017 |
| biolink:decreases_response_to | 3,367 |
| biolink:has_metabolite | 3,048 |
| biolink:increases_response_to | 2,936 |
| biolink:disease_has_basis_in | 2,799 |
| biolink:associated_with_increased_likelihood_of | 2,561 |
| biolink:has_substrate | 2,095 |
| biolink:composed_primarily_of | 1,725 |
| biolink:has_plasma_membrane_part | 1,608 |
| biolink:negatively_correlated_with | 1,467 |
| biolink:exacerbates_condition | 1,428 |
| biolink:is_frameshift_variant_of | 1,035 |

### S4. Edge types (continued)

| Edge type (Biolink predicate) | Edge count |
|---|---|
| biolink:is_input_of | 876 |
| biolink:temporally_related_to | 841 |
| biolink:assesses | 831 |
| biolink:narrow_match | 790 |
| biolink:is_nonsense_variant_of | 671 |
| biolink:mentions | 535 |
| biolink:opposite_of | 473 |
| biolink:ameliorates_condition | 396 |
| biolink:acts_upstream_of_or_within_positive_effect | 370 |
| biolink:acts_upstream_of_positive_effect | 365 |
| biolink:has_increased_amount | 262 |
| biolink:has_decreased_amount | 230 |
| biolink:acts_upstream_of_negative_effect | 162 |
| biolink:acts_upstream_of_or_within_negative_effect | 142 |
| biolink:has_molecular_consequence | 74 |
| biolink:has_not_completed | 65 |
| biolink:in_preclinical_trials_for | 60 |
| biolink:model_of | 39 |
| biolink:broad_match | 34 |
| biolink:has_chemical_role | 24 |
| biolink:homologous_to | 21 |
| biolink:indirectly_physically_interacts_with | 2 |
| biolink:in_complex_with | 2 |
| biolink:beneficial_in_models_for | 1 |
| biolink:enabled_by | 1 |

### S5. Example node and edge records

Each line of KRAKEN's nodes and edges NDJSON files is one JSON record. The records below are the node and edge shown in Figure 1, reproduced from the files and indented here for readability (in the files each record occupies a single line).

#### Example node record


```
{
  "id": "RM:0028430",
  "categories": [
    "biolink:SmallMolecule"
  ],
  "provided_by": [
    "infores:loinc",
    "infores:multiomics-clinicaltrials",
    "infores:refmet",
    "infores:robokop-kg",
    "infores:rtx-kg2",
    "infores:sri-node-normalizer",
    "infores:umls-metathesaurus",
    "translator-kg-open"
  ],
  "equivalent_ids": [
    "CAS:200-466-7",
    "CAS:60-27-5",
    "CAS:82016-55-5",
    "CHEBI:14029",
    "CHEBI:16737",
    "CHEBI:23406",
    "CHEBI:3910",
    "CHEMBL.COMPOUND:CHEMBL4849635",
    "CHEMBL.COMPOUND:CHEMBL65567",
    "CHV:0000003442",
    "DRUGBANK:DB11846",
    "HMDB:HMDB0000562",
    "INCHIKEY:DDRJAANPRJIHGJ-UHFFFAOYSA-N",
    "KEGG.COMPOUND:C00791",
    "KEGG:C00791",
    "LOINC:LP14355-9",
    "LOINC:LP32035-5",
    "MESH:D003404",
    "NCIT:C399",
    "PUBCHEM.COMPOUND:137319715",
    "PUBCHEM.COMPOUND:588",
    "REACT:R-ALL-30733",
    "REACT:R-ALL-446592",
    "RM:0028430",
    "RXCUI:2913",
    "RXNORM:2913",
    "SMILES:CN1CC(=O)NC1=N",
    "SNOMEDCT:15373003",
    "UMLS:C0010294",
    "UNII:AYI8EX34EU"
  ],
  "name": "Creatinine",
  "description": "A lactam obtained by formal cyclocondensation of creatine. It is a metabolite of creatine.",
  "synonyms": [
    "1-methylglycocyamidine",
    "1-methylhydantoin-2-imide",
    "2-Imino-1-methyl-4-imidazolidinone",
    "CREAT [cytosol]",
    "CREAT [extracellular region]",
    "CREATININE",
    "Creatinine (substance)",
    "Creatinine [divisor]",
    "Kreatininum",
    "NSC-13123",
```

### S5. Example node and edge records (continued)


```
    "creatinine"
  ],
  "urls": [
    "http://purl.obolibrary.org/obo/CHEBI_16737",
    "https://identifiers.org/reactome:R-ALL-30733",
    "https://identifiers.org/reactome:R-ALL-446592",
    "https://www.metabolomicsworkbench.org/databases/refmet/refmet_details.php?REFMET_ID=RM0028430"
  ],
  "chemical_formula": "C4H7N3O",
  "exact_mass": 113.058912,
  "attributes": {
    "infores:refmet": {
      "main_class": "Azoles",
      "super_class": "Organoheterocyclic compounds",
      "sub_class": "Imidazolines"
    },
    "infores:robokop-kg": {
      "CHEBI_ROLE_human_metabolite": true,
      "CHEBI_ROLE_metabolite": true,
      "CHEBI_ROLE_biochemical_role": true,
      "CHEBI_ROLE_diagnostic_agent": true,
      "CHEBI_ROLE_mammalian_metabolite": true,
      "CHEBI_ROLE_pharmaceutical": true,
      "CHEBI_ROLE_eukaryotic_metabolite": true,
      "CHEBI_ROLE_animal_metabolite": true,
      "information_content": 76.6
    },
    "translator-kg-open": {
      "xref": [
        "InChIKey:DDRJAANPRJIHGJ-UHFFFAOYSA-N",
        "SMILES:CN1CC(=O)NC1=N"
      ],
      "chembl_natural_product": true,
      "information_content": 76.5
    },
    "infores:sri-node-normalizer": {
      "information_content": 76.78008365410358
    },
    "infores:rtx-kg2": {
      "canonical_category": "biolink:SmallMolecule"
    },
    "infores:loinc": {
      "PartTypeName": "COMPONENT",
      "Status": "ACTIVE"
    }
  }
}
```

### S5. Example node and edge records (continued)

#### Example edge record


```
{
  "subject": "RM:0028430",
  "object": "MONDO:0005575",
  "predicate": "biolink:correlated_with",
  "primary_knowledge_source": "infores:hmdb",
  "knowledge_level": "knowledge_assertion",
  "agent_type": "manual_agent",
  "aggregator_knowledge_source": [
    "infores:robokop-kg"
  ],
  "attributes": {
    "infores:robokop-kg": {
      "original_subject": "HMDB:HMDB0000562",
      "original_object": "OMIM:114500"
    }
  }
}
```

### S6. Figure queries

Links open the query in the KRAKEN web interface (https://app.krakenkg.com). Results reflect KRAKEN v2.3.0 and may change in later releases. Figures 5, 6 and 7 were generated from the v2.3.0 graph files with the plotting scripts in the KRAKEN repository (scripts/visualization) and have no corresponding interface query.

#### Figure 1A

Node example. Looks up Creatinine (RM:0028430) by identifier, which opens its Node Details panel directly.

`https://app.krakenkg.com/#/search?ids=RM:0028430`

#### Figure 1B

Edge example. Run the query, expand the first result, select the 'correlated with' edge between Creatinine and colorectal cancer, and choose 'Show attributes'.

`https://app.krakenkg.com/#/path?start=RM:0028430&end=MONDO:0005575&hops=1-2&ranking=established&traversal=deep_dive`

#### Figure 2

Multi-hop query for pathways and biological processes connected to cholesterol through proteins. Select the end node of the first result to open its Node Details panel.

`https://app.krakenkg.com/#/path?start=CHEBI:16113&end_category=Pathway,BiologicalProcessOrActivity&hops=1-2&ranking=established&traversal=deep_dive&intermediate_node_category.in=biolink:Protein`

#### Figure 3

Query builder for the Figure 2 query. Open the Figure 2 link and choose 'Edit Query'.

`https://app.krakenkg.com/#/path?start=CHEBI:16113&end_category=Pathway,BiologicalProcessOrActivity&hops=1-2&ranking=established&traversal=deep_dive&intermediate_node_category.in=biolink:Protein`

#### Figure 4

Node search for 'atrial fibrillation'. Select the first result to open its Node Details panel.

`https://app.krakenkg.com/#/search?text=atrial%20fibrillation`

#### Figure 8

Enrichment analysis of the 88 signature proteins against pathways and biological processes, restricted to participation edges (default 1,000 permutations). To list all significant targets, change limit=30 to limit=500.

`https://app.krakenkg.com/#/enrichment?nodes=NCBIGene:147945,NCBIGene:3952,NCBIGene:133,NCBIGene:3484,NCBIGene:2167,NCBIGene:177,NCBIGene:1435,NCBIGene:3603,NCBIGene:92737,NCBIGene:5653,NCBIGene:56477,NCBIGene:10673,NCBIGene:6441,NCBIGene:54,NCBIGene:181,NCBIGene:1956,NCBIGene:117156,NCBIGene:4319,NCBIGene:4973,NCBIGene:4151,NCBIGene:5806,NCBIGene:8788,NCBIGene:2277,NCBIGene:9965,NCBIGene:3627,NCBIGene:3977,NCBIGene:1476,NCBIGene:8797,NCBIGene:920,NCBIGene:3558,NCBIGene:1473,NCBIGene:6401,NCBIGene:58191,NCBIGene:923,NCBIGene:27189,NCBIGene:4854,NCBIGene:8685,NCBIGene:2658,NCBIGene:6373,NCBIGene:7037,NCBIGene:1357,NCBIGene:11093,NCBIGene:3958,NCBIGene:3596,NCBIGene:50604,NCBIGene:763,NCBIGene:5328,NCBIGene:7039,NCBIGene:4353,NCBIGene:9048,NCBIGene:6504,NCBIGene:3566,NCBIGene:4313,NCBIGene:26291,NCBIGene:83886,NCBIGene:7079,NCBIGene:56729,NCBIGene:127482540,NCBIGene:142,NCBIGene:355,NCBIGene:6370,NCBIGene:3593,NCBIGene:3976,NCBIGene:145264,NCBIGene:3339,NCBIGene:1522,NCBIGene:8600,NCBIGene:3425,NCBIGene:8743,NCBIGene:2250,NCBIGene:100,NCBIGene:9332,NCBIGene:53832,NCBIGene:5327,NCBIGene:3605,NCBIGene:90865,NCBIGene:23584,NCBIGene:126014,NCBIGene:57823,NCBIGene:8993,NCBIGene:133395145,NCBIGene:4803,NCBIGene:6347,NCBIGene:654,NCBIGene:23765,NCBIGene:51744,NCBIGene:6359,NCBIGene:3559&target=Pathway,BiologicalProcessOrActivity&predicate=has_participant,participates_in&limit=30`

#### Figure 9

Subgraph query over the 41 measured inflammatory and immune proteins. Select 'cytokine-mediated signaling pathway' in the Intermediate Nodes table to highlight it.

`https://app.krakenkg.com/#/subgraph?nodes=NCBIGene:9048,NCBIGene:6359,NCBIGene:6347,NCBIGene:6370,NCBIGene:56477,NCBIGene:9332,NCBIGene:51744,NCBIGene:920,NCBIGene:923,NCBIGene:1401,NCBIGene:1435,NCBIGene:3627,NCBIGene:6373,NCBIGene:58191,NCBIGene:355,NCBIGene:3593,NCBIGene:3596,NCBIGene:3603,NCBIGene:3605,NCBIGene:27189,NCBIGene:23765,NCBIGene:3558,NCBIGene:50604,NCBIGene:53832,NCBIGene:3559,NCBIGene:90865,NCBIGene:3566,NCBIGene:3958,NCBIGene:3976,NCBIGene:3977,NCBIGene:8685,NCBIGene:4353,NCBIGene:4973,NCBIGene:8993,NCBIGene:5806,NCBIGene:6504,NCBIGene:57823,NCBIGene:8797,NCBIGene:8743,NCBIGene:8600,NCBIGene:10673&hops=2&limit=50&ranking=deep_dive&traversal=deep_dive&intermediate_node_category.in=biolink:Protein,biolink:Gene,biolink:Pathway&prefix.in=NCBIGene,HGNC,GO,REACT&agent_type.not_in=text_mining_agent`

### S6. Figure queries (continued)

#### Figure 10A

Query graph: IL5RA, mepolizumab (with equivalent terms), asthma. Text-mined edges excluded.

https://app.krakenkg.com/#/querygraph?n0=ATC:R03DX09&n0.category=Drug,Protein&n0.expand=exact_match,close_match,same_as&n0.expand.hops=1&n0.expand.agent_type.not_in=text_mining_agent&n1=HGNC:6017&n2=MONDO:0004979&e0=n0-n1&e0.agent_type.not_in=text_mining_agent&e1=n0-n2&e1.predicate=treats_or_applied_or_studied_to_treat&e1.agent_type.not_in=text_mining_agent&ranking=established&traversal=deep_dive&layout=n0.757.91,n1.557.91,n2.1057.91

#### Figure 10B

Query graph: MMP10, marimastat (with equivalent terms), asthma (with subtypes, parent types and equivalent terms, up to two hops). Text-mined edges excluded.

https://app.krakenkg.com/#/querygraph?n0=RM:0224430&n0.category=Drug,SmallMolecule&n0.expand=exact_match,close_match,same_as&n0.expand.hops=1&n0.expand.agent_type.not_in=text_mining_agent&n1=HGNC:7156&n2=MONDO:0004979&n2.expand=subclass_of,superclass_of,exact_match,close_match,same_as&n2.expand.hops=2&n2.expand.agent_type.not_in=text_mining_agent&e0=n0-n1&e0.agent_type.not_in=text_mining_agent&e1=n0-n2&e1.predicate=treats_or_applied_or_studied_to_treat&e1.agent_type.not_in=text_mining_agent&paths_per_pair=1&ranking=established&traversal=deep_dive&layout=n0.757.91,n1.557.91,n2.1057.91

#### Figure 10C

Query graph: TNFRSF8 (CD30), brentuximab vedotin (Adcetris; with equivalent terms), asthma (with subtypes, parent types and equivalent terms, up to two hops). Text-mined edges excluded.

https://app.krakenkg.com/#/querygraph?n0=ATC:L01FX05&n0.category=Drug,SmallMolecule,Protein&n0.expand=exact_match,close_match,same_as&n0.expand.hops=1&n0.expand.agent_type.not_in=text_mining_agent&n1=HGNC:11923&n2=MONDO:0004979&n2.expand=subclass_of,superclass_of,exact_match,close_match,same_as&n2.expand.hops=2&n2.expand.agent_type.not_in=text_mining_agent&e0=n0-n1&e0.agent_type.not_in=text_mining_agent&e1=n0-n2&e1.predicate=treats_or_applied_or_studied_to_treat&e1.agent_type.not_in=text_mining_agent&paths_per_pair=2&ranking=established&traversal=deep_dive&layout=n0.757.91,n1.557.91,n2.1057.91
